# NAP1 switches from an activator to a limiter of interferon induction by trapping TBK1 in condensates

**DOI:** 10.1101/2025.05.21.655319

**Authors:** Damien Glon, Quentin Riller, Frédéric Rivière, Benjamin Léonardon, Ariane Guillemot, Laïla Sago, Olivier Pellé, Duong Ho-Nhat, Karine Brochard, Brigitte Bader-Meunier, Marie-Louise Frémond, Alice Lepelley, Yanick Crow, Maud Tusseau, Alexandre Belot, Cécile Lagaudrière-Gesbert, Frédéric Rieux-Laucat, Yves Gaudin

## Abstract

TBK1 kinase is a central regulator of type I IFN production. Upon activation of the IFN-β induction pathway, TBK1-adaptor proteins (NAP1, SINTBAD, TANK) form condensates with liquid properties. We showed that NAP1 condensates concentrate TBK1. Using NAP1^KO^ cell lines, we discovered that NAP1 exerts a dual effect on TBK1 activity. Initially, NAP1 binds TBK1 and increases its activity, which enhances the activation of the IFN pathway. Then, phosphorylation of NAP1 by TBK1 induces the formation of NAP1 condensates. These condensates concentrate TBK1 and PP2A, a phosphatase known to dephosphorylate and consequently deactivate TBK1, thus limiting IFN induction. Additionally, in patients with lupus or interferonopathies, we identified NAP1 variants, unable to form condensates upon cell exposure to danger signals, which can only activate TBK1 without limiting its activity. This study reveals an original mode of regulating a signaling pathway by formation of condensates and provides a molecular explanation for certain interferonopathies.

## Introduction

Type I interferons (IFNs) exhibit potent antiviral, antiproliferative and immunomodulatory effects (1). They are expressed and secreted following pathogen-associated molecular patterns (PAMPs) recognition by dedicated cellular proteins known as pattern recognition receptors (PRRs) (2). PRRs include Toll-like receptors (TLRs), which are transmembrane proteins recognizing extracellular PAMPs from several pathogens(3–5), the cytoplasmic RIG-I-like receptors (RLRs) involved in the recognition of RNA viruses (by sensing cytoplasmic dsRNA) (6), and the cyclic GMP-AMP synthase (cGAS) that detect the presence of dsDNA in the cytosol of eukaryotic cells (7).

TANK-binding kinase 1 (TBK1) is a central regulator of type I IFN production (8–10). It is located at the crossroads of several signaling pathways, including those triggered by TLR3, RLRs, and cGAS (10,11). Indeed, the binding of dsRNA by TLR3 induces a signaling cascade that leads to the dimerization and autophosphorylation at Ser172 of TBK1 (12), two essential features for kinase activation. Similarly, recognition of dsRNA by RLRs (like RIG-I or MDA5) results in their association with the mitochondrial antiviral signaling (MAVS) protein at the outer mitochondrial membrane, which induces MAVS oligomerization and assembly of TBK1-activating complexes (13). Finally, cytosolic dsDNA binding stimulates cGAS enzymatic activity (11). The generated cyclic dinucleotide 2′,3′-cGAMP then acts as a second messenger by binding and activating the stimulator of IFN genes (STING). Upon activation, STING undergoes conformational changes that allow its translocation from the endoplasmic reticulum to the Golgi apparatus complex and the recruitment of TBK1 (11). Once activated, TBK1 phosphorylates transcription factors IRF3 and/or IRF7 (8,9), resulting in their homo- or heterodimerization and subsequent import into the nucleus to induce expression of type I IFN genes.

Subsequent binding of IFNs to their receptors leads to the upregulation of hundreds of IFN-stimulated genes (ISGs) that establish an antiviral state (14). Because of the large number of genes activated by type I IFNs, their production must be finely regulated. Indeed, uncontrolled IFN induction in the context of genetic diseases leads to interferonopathies and systemic lupus erythematosus (SLE) (15,16). TKB1 kinase regulation is a crucial element for the precise orchestration of these pathways and TBK1 activity is controlled by several mechanisms that are not yet fully identified (17).

Among TBK1-adaptor proteins, NAP1 (encoded by the *AZI2* gene) (18,19), SINTBAD/TBKBP1 (20) and TANK (21) are homologous proteins sharing the same modular organization (Figure S1)(22). They possess a central TBK1 binding domain (TBD), made of a single α-helix, which associates with the C-terminal adaptor binding domain (ABD) of TBK1 as demonstrated for NAP1 TBD (23). The TBD is flanked on its N-terminal side by a homodimerization domain (HD), which contains long α-helices and intrinsically disordered domains (IDDs). The C-terminal domain (CTD) of the adaptor proteins is also largely disordered and, in the case of SINTBAD and TANK, contains a zinc finger (ZF) (Figure S1).

These adaptor proteins compete for TBK1 binding on the C-terminal kinase domain (ABD, adaptor binding domain) (20,24). Their role in type I IFN induction remains controversial. Some loss-of-function RNAi experiments indicate that they are required for type I IFN production in response to danger signals (19,20,25). Moreover, their ectopic expression leads to increased IFN production(18–20,25). However, this view was challenged by the recent observation that TANK-deficient mice have no apparent defect in type I IFN production (26) and that the triple deficiency in TBK1-adaptor proteins does not impair the RLRs-MAVS dependent pathway (27). Recently, our group showed that in the presence of various danger signals, NAP1 and SINTBAD form condensates with liquid properties (28), potentially providing a new level of regulation of the IFN induction pathway.

In this study, we characterized the condensates formed by the three TBK1-adaptor proteins. We confirmed their liquid properties and identified the domains involved in their formation. Our results indicated that TBK1 is recruited into NAP1 condensates but interferes with the formation of SINTBAD and TANK condensates. We also identified the protein domains involved in these processes. Importantly, we discovered that NAP1 actually plays a dual role on the TBK1 kinase activity. First, by binding TBK1, NAP1 increases its activity and therefore leads to enhanced activation of the IFN pathway. In parallel, NAP1 is phosphorylated by TBK1. This phosphorylation lowers the critical concentration of NAP1 favoring the formation of condensates concentrating TBK1 and the phosphatase complex PP2A. Within these condensates, the PP2A complex dephosphorylates TBK1, which limits the IFN activation pathway. Finally, we identified NAP1 variants in patients with SLE or interferonopathies that are unable to form condensates upon exposure of cells to danger signals and are therefore only able to activate TBK1 but not limit its activity.

Our work highlights a novel and original way of regulating the activity of an enzyme by the formation of liquid condensates and provides a pathophysiological mechanism in patients with interferonopathies or lupus.

## Results

### Endogenous TBK1, NAP1, SINTBAD and TANK are found in cytoplasmic condensates upon exposure to danger signals

The ability of TBK1 and its adaptor proteins NAP1/AZI2 and SINTBAD/TBK1BP1 to form or be recruited into condensates upon exposure of the cell to several distinct stresses was already documented. Here, using specific antibodies and immunofluorescence (IF), we confirmed that endogenous TBK1, NAP1, SINTBAD and additionally TANK were found in cytoplasmic condensates in both HEK293T and HAP1 cell lines upon infection by rabies virus (RABV) (Figures 1A and S2A) or after transfection with 5’ppp-dsRNA (Figures 1B and S2C).

**Fig. 1.**
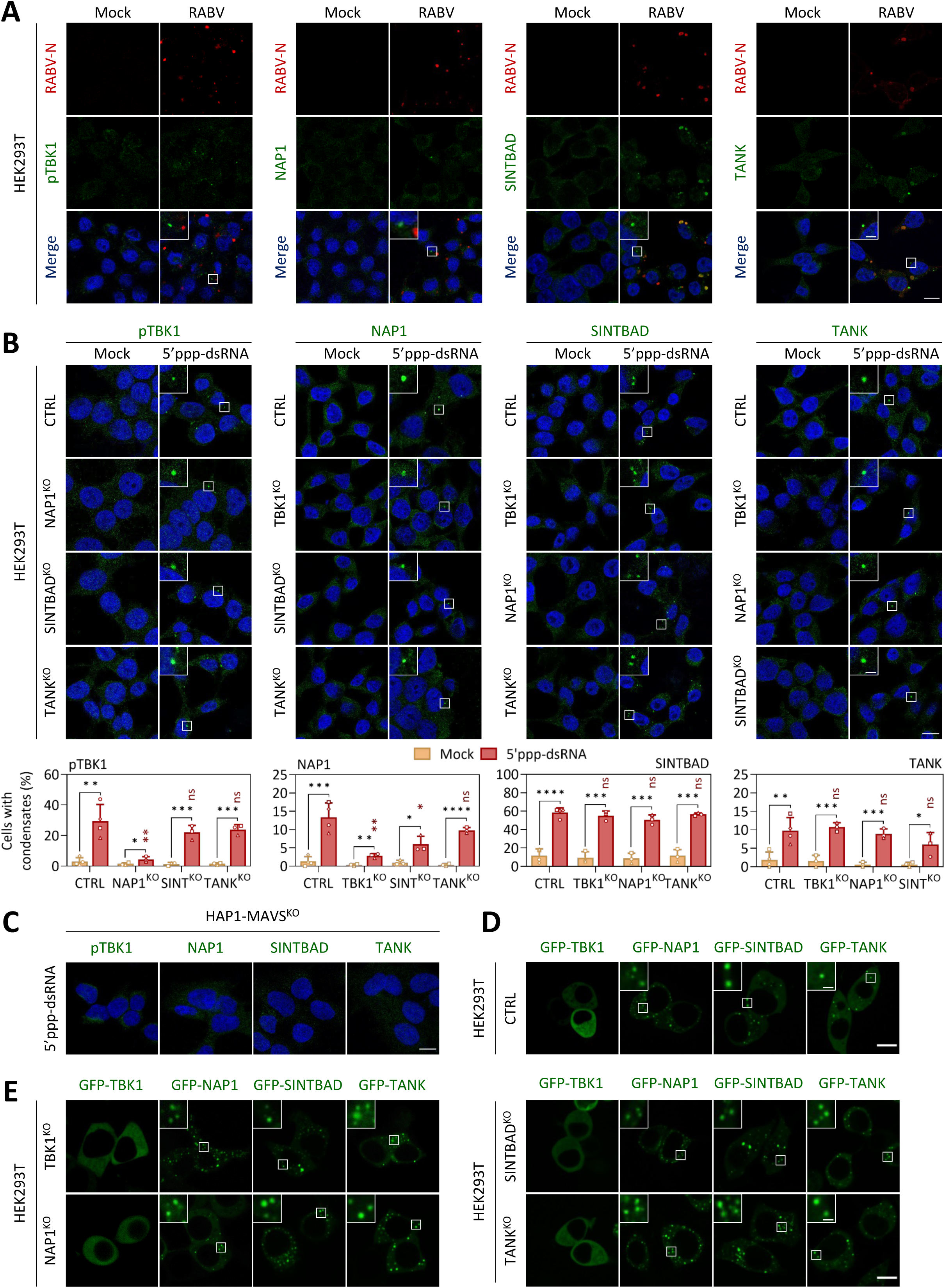
Detection of danger signals as well as ectopic expression of TBK1, NAP1, SINTBAD, or TANK induce their condensation. **A)** HEK293T cells were infected with RABV (MOI=3) or not (Mock). At 16 h p.i., cells were fixed before co-staining with anti-RABV-N and either anti-pTBK1, anti-NAP1, anti-SINTBAD, or anti-TANK antibodies and analyzed by confocal microscopy. **B)** Parental (CTRL) and KO HEK293T cells (TBK1^KO^, NAP1^KO^, SINTBAD^KO^ and TANK^KO^) were incubated with a medium containing LyoVec alone (Mock) or transfected with 5’ppp-dsRNA using LyoVec (5’ppp-dsRNA). After 6 h, cells were fixed before staining with anti-pTBK1, anti-NAP1, anti-SINTBAD, or anti-TANK antibody and analyzed by confocal microscopy. The percentage of cells with condensates is indicated under the corresponding images. The bars represent the mean ± SD of at least 3 independent experiments and the corresponding data points are shown. Statistical significance between cells treated or not by 5’ppp-dsRNA is indicated (in black). Statistical significance between control and KO cells treated by 5’ppp-dsRNA is indicated (in red). ns =non-significant; *P < 0.05 ; **P < 0.01 ; ***P < 0.001 ; ****P < 0.0001 (Student t-test). **C)** MAVS^KO^ HAP1 cells were transfected with 5’ppp-dsRNA using Lipofectamine 3000 (5’ppp-dsRNA). After 6 h, cells were fixed before staining with anti-pTBK1, anti-NAP1, anti-SINTBAD or anti-TANK antibody and analyzed by confocal microscopy. **D-E)** Parental (CTRL) (**D**) and KO HEK293T cells (TBK1^KO^, NAP1^KO^, SINTBAD^KO^ and TANK^KO^ (**E**) were transfected with plasmids encoding the indicated GFP versions of TBK1, NAP1, SINTBAD or TANK. After 20 h, cells were analyzed by confocal microscopy. On microscopy images, square boxes have been magnified (3X) at the top left. Scale bars: 10µm and 2 µm for insets.

We then investigated the requirement of TBK1 or its adaptor proteins in the formation of those condensates. For this, we used TBK1^KO^, NAP1^KO^, SINTBAD^KO^ and TANK^KO^ HAP1 (that are commercially available) or HEK293T (that we constructed) cell lines (Figure S2B). In NAP1^KO^, TANK^KO^ or SINTBAD^KO^ cell lines, endogenous TBK1 condensates were observed upon cell transfection with 5’ppp-dsRNA, indicating that no adaptor proteins are mandatory for their formation (Figures 1B and S2C). However, significantly fewer NAP1^KO^ HEK293T cells than the parental cell line contained TBK1 condensates, suggesting that NAP1 plays a key role in TBK1 condensation (Figure 1B).

Upon cell transfection with 5’ppp-dsRNA, NAP1 condensates were observed in TBK1^KO^, SINTBAD^KO^ or TANK^KO^ cell lines (Figures 1B and S2C). However, significantly fewer TBK1^KO^ and SINTBAD^KO^ HEK293T cells than the parental cell line contained endogenous NAP1 condensates, suggesting that TBK1 and SINTBAD could regulate NAP1 condensation (Figure 1B).

SINTBAD and TANK condensates were observed in all knock-out (KO) cell lines (Figures 1B and S2C) with the same proportion of cells containing condensates compared to the parental cell line (Figure 1B), indicating that neither another adaptor protein nor TBK1 is necessary for their formation (Figures 1B and S2C). Notably, no adaptor protein condensates were observed when MAVS^KO^ HAP1 cells (Figure S2B) were transfected with 5’ppp-dsRNA, demonstrating that their formation depends on the RLR pathway (Figure 1C).

Since type I IFN can also be induced through the cGAS-STING pathway, we investigated whether TBK1 condensates are present in cells treated with diABZI, a STING agonist. For this purpose, we used THP-1 cells known to express STING, unlike HEK293T and HAP1 cells. When THP-1 cells were exposed to diABZI for 6h, TBK1 condensation was observed (Figure S2D). We also transfected HEK293T cells with a plasmid encoding STING. One day after transfection, we treated the cells with diABZI and observed that TBK1 formed condensates only in STING-expressing HEK293T cells (Figure S2E). These results demonstrated that TBK1 condensation is induced by activation of both RLR-MAVS and cGAS-STING pathways.

### Ectopic TBK1-adaptor proteins expression leads to condensates formation

We previously showed that ectopic expression of fluorescent versions of NAP1 and SINTBAD (either fused to GFP or mCherry [mC] at their N-terminus) leads to the formation of condensates of these proteins (28). We transfected HEK293T cells with pcDNA plasmids encoding fluorescent versions of the adaptor proteins. We confirmed our previous observations and showed that TANK also formed condensates when ectopically expressed (Figure 1D). On the other hand, cell transfection with plasmid encoding GFP-TBK1 did not result in the formation of condensates of this latter protein (Figure 1D). The same observations were made in the different KO HEK293T cells, revealing that the formation of condensates by ectopically expressed adaptor proteins does not depend on the presence of another endogenous adaptor nor on the presence of TBK1 (Figure 1E). It is noteworthy that GFP-TBK1 condensation was observed when HEK293T cells were exposed to danger signals, such as transfection with 5’ppp-dsRNA or infection by RABV (Figures S2F-G).

Importantly, the formation of fluorescent adaptor protein condensates was not due to the fusion with GFP or mC proteins since the condensates were also observed by IF when cells were transfected with plasmids, allowing the ectopic expression of the untagged adaptor proteins (Figure S2H).

### Condensates formed by TBK1-adaptor proteins have liquid properties

We then characterized the dynamics of the condensates formed by the GFP-tagged adaptor proteins and assessed their response to various treatments by live imaging. First, we demonstrated that the adaptor protein condensates disappeared within seconds when transfected cells were exposed to 10% of 1,6-hexanediol (1,6-HD - Figure 2A). They were also sensitive to cellular hypotonic shock, a hallmark of many biomolecular condensates formed by liquid liquid phase separation (LLPS) (29)(Figure 2B). After the hypotonic shock, during the return to cellular homeostasis, the condensates gradually reappeared within a few minutes (Figure 2B). In addition, we also observed some fusion events between these condensates (Figure 2C), further supporting their liquid-like properties.

**Fig. 2.**
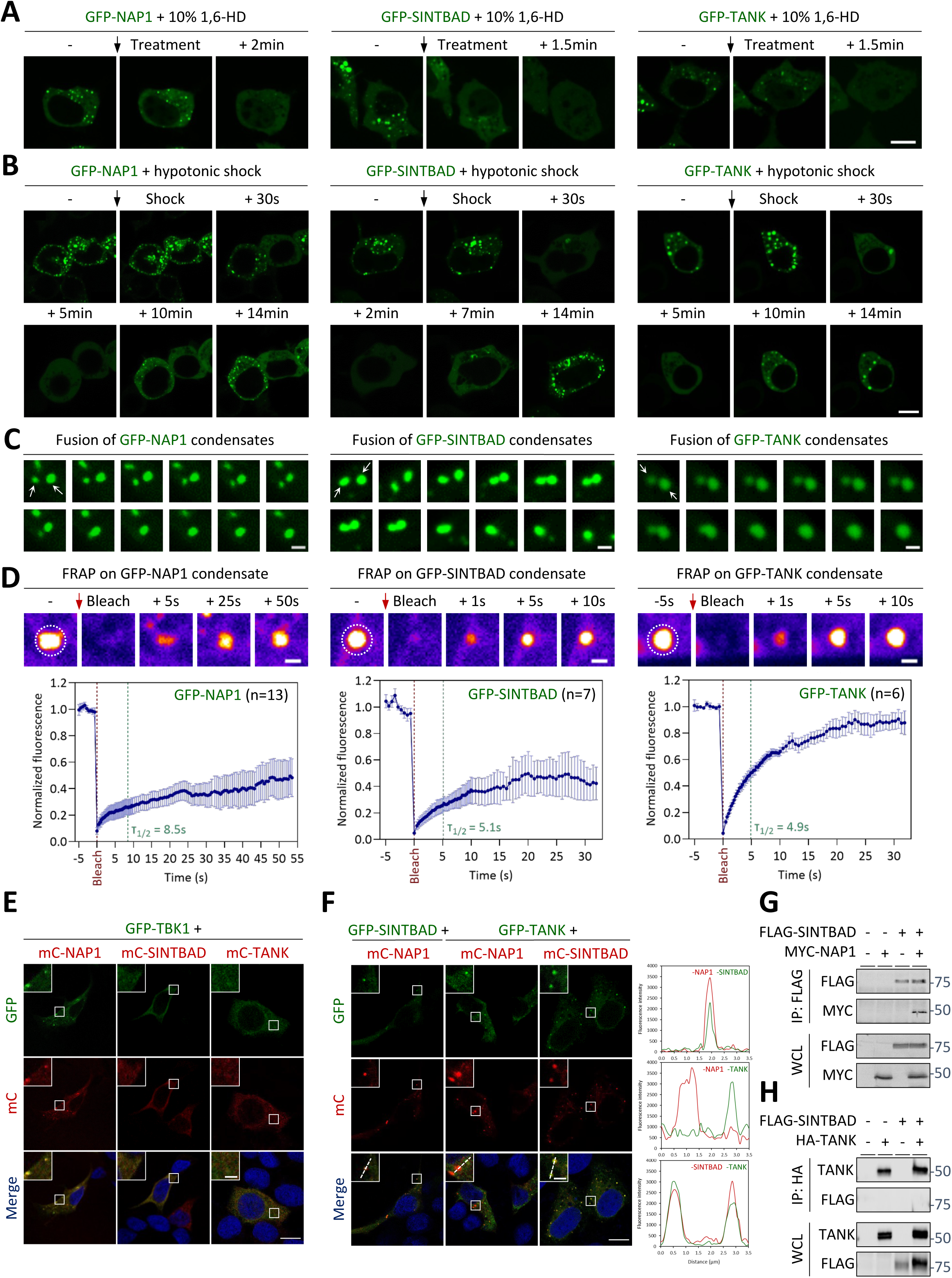
Liquid properties of NAP1, SINTBAD and TANK condensates and their interplay with TBK1. **A-B)** HEK293T cells were transfected with plasmids encoding GFP-NAP1, GFP-SINTBAD or GFP-TANK. After 20 h, live cells were analyzed by a confocal spinning-disk microscope and treated with 10% of 1,6-HD **(A)** or submitted to a hypotonic shock **(B)**. **C)** HEK293T cells were transfected with plasmids encoding GFP-NAP1, GFP-SINTBAD or GFP-TANK. After 20 h, live cells were analyzed by a confocal spinning-disk microscope. Fusions between condensates are indicated by arrows. Images were taken every 500 ms. **D)** HEK293T cells were transfected with plasmids encoding GFP-NAP1, GFP-SINTBAD or GFP-TANK. After 20 h, whole condensates were photobleached and the recovery of fluorescence was followed by a confocal spinning-disk microscope. A representative FRAP experiment is shown for each protein and the photobleached region (Ø=3 µm) is indicated. For each protein, the mean curve and the SD of the indicated number of FRAP experiments is shown. **E)** HAP1 cells were co-transfected with plasmids encoding GFP-TBK1 and either mC-NAP1, mC-SINTBAD, or mC-TANK. After 24 h, cells were fixed and analyzed by confocal microscopy. **F)** HAP1 cells were co-transfected with plasmids encoding the indicated fluorescent versions of TBK1-adaptor proteins. After 24 h, cells were fixed and analyzed by confocal microscopy. The fluorescence corresponding to each protein along the lines traced over condensates is indicated on the graphs. **G)** HEK293T cells were co-transfected with plasmids encoding FLAG-SINTBAD and MYC-NAP1 for 24 h. After immunoprecipitation of SINTBAD with an anti-FLAG antibody, cell lysates (WCL) and immunoprecipitated proteins (IP) were analyzed by WB with anti-FLAG and anti-MYC antibodies. **H)** HEK293T cells were co-transfected with plasmids encoding FLAG-SINTBAD and HA-TANK for 24 h. After immunoprecipitation of TANK with an anti-HA antibody, cell lysates (WCL) and immunoprecipitated proteins (IP) were analyzed by WB with anti-TANK and anti-SINTBAD antibodies. On microscopy images, square boxes have been magnified (3X) at the top left. Scale bars: 10µm and 2 µm for insets and FRAP experiments.

To investigate the dynamics of these condensates, we performed fluorescence recovery after photobleaching (FRAP) experiments on condensates formed by GFP-NAP1, GFP-SINTBAD, or GFP-TANK (Figure 2D). Whole condensates were photobleached. The fluorescence signal recovery was complete and fast for GFP-TANK (t_1/2_∼5s), whereas it was partial for NAP1 and SINTBAD with a mobile fraction of approximately 40%. This indicated that adaptor proteins could shuttle between their cytoplasmic pool and the condensates. Collectively, these experiments, as well as the spherical morphology of the condensates, were consistent with their formation by LLPS.

### TBK1 accumulates inside NAP1 condensates but disrupts SINTBAD and TANK condensates

To investigate the interplay between TBK1 and its adaptor proteins, we co-transfected HAP1 cells with plasmids encoding two distinct proteins, one fused to GFP, and the other to mC. As previously shown (28), the co-expression of GFP-TBK1 and mC-NAP1 induced the recruitment of TBK1 into NAP1 condensates (Figure 2E). This contrasted with the absence of condensates in cells co-expressing GFP-TBK1 and mC-SINTBAD or mC-TANK, indicating that TBK1 negatively affects SINTBAD and TANK condensate formation (Figure 2E).

Next, we investigated the interplay between adaptor proteins. Co-expression of the fluorescent versions of SINTBAD and NAP1 (or TANK) revealed that SINTBAD condensates colocalized with those formed by NAP1 (or TANK). However, the co-expression of fluorescent versions of NAP1 and TANK revealed that they formed distinct condensates (Figure 2F).

The colocalization of the condensates formed by SINTBAD with those formed by NAP1 and by TANK suggested that SINTBAD could interact with the two other adaptor proteins. To test this, we co-transfected a FLAG-tagged version of SINTBAD with either a MYC-tagged version of NAP1 or an HA-tagged version of TANK. SINTBAD was co-immunoprecipitated with NAP1 (Figure 2G), while this was not the case with TANK (Figure 2H). This suggested that the colocalization of SINTBAD and TANK condensates did not imply a strong interaction between these two proteins.

### The homodimerization domain of TBK1-adaptor proteins is sufficient for condensate formation

We took advantage of this expression/co-expression system to identify the molecular bases underlying the formation of condensates by TBK1-adaptor proteins, the recruitment of TBK1 into NAP1 condensates, as well as the negative impact of TBK1 on SINTBAD and TANK condensates.

For each TBK1-adaptor protein, we constructed two deletion mutants (fused at their N-terminus to mC or GFP) corresponding to the HD (residues 1-211 for NAP1, 1-280 for SINTBAD and 1-129 for TANK) and the TBD plus CTD (TBD-CTD, residues 210-392 for NAP1, 280-615 for SINTBAD and 129-425 for TANK) (Figures S1 and 3A). The transfection of cells by plasmids expressing these deletion mutants revealed that, for each adaptor protein, the HD was sufficient to observe condensates. On the contrary, the expression of TBD-CTD alone did not induce condensate formation (Figure 3B).

**Fig. 3.**
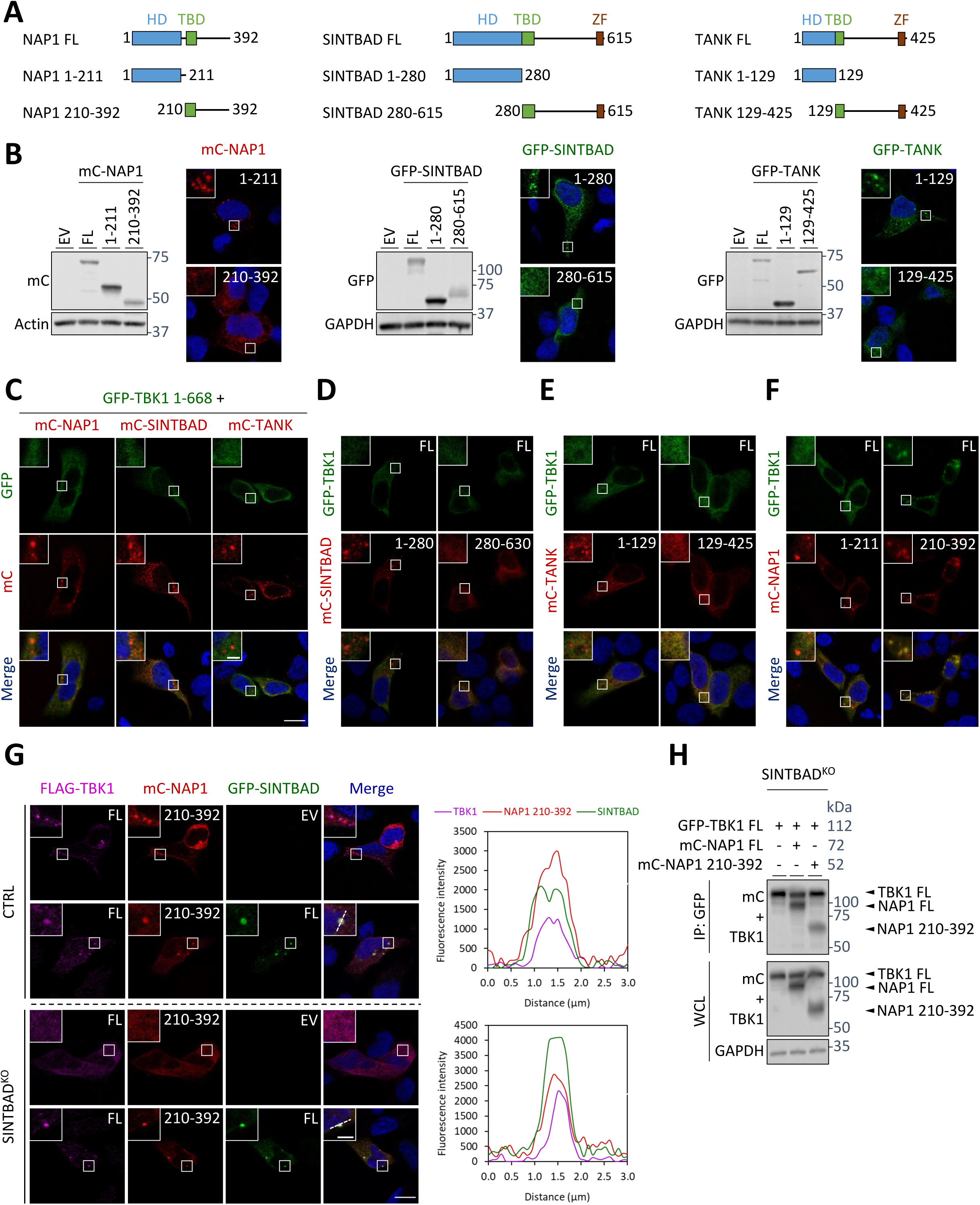
The HD of adaptor proteins is sufficient to induce condensate formation and TBK1 ABD is required for its recruitment into NAP1 condensates and to inhibit SINTBAD and TANK condensates formation. **A)** Domain organization of NAP1, SINTBAD and TANK and their deletion mutants used in this study. HD, homodimerization domain; TBD, TBK1 binding domain; ZF, zinc finger. **B)** HEK293T cells were transfected with plasmids encoding NAP1, SINTBAD and TANK truncations fused to an mC- or a GFP-tag in their N-terminal part. After 24 h, cells were either lysed for analysis of truncated adaptor proteins expression by WB (using anti-mC or anti-GFP and anti-Actin or anti-GAPDH antibodies) or fixed for analysis by confocal microscopy. **C)** HAP1 cells were co-transfected with plasmids encoding GFP-TBK1 1-668 and either mC-NAP1, mC-SINTBAD or mC-TANK. After 24 h, cells were fixed and analyzed by confocal microscopy. **D-F)** HAP1 cells were co-transfected with plasmids encoding GFP-TBK1 FL and indicated truncations of mC-SINTBAD (**D**), mC-TANK (**E**) or mC-NAP1 (**F**). After 24 h, cells were fixed and analyzed by confocal microscopy. **G)** Parental (CTRL) and SINTBAD^KO^ HAP1 cells were co-transfected with plasmids encoding FLAG-TBK1 FL, mC-NAP1 210-392 and GFP-SINTBAD FL or an empty vector (EV). After 24 h, cells were fixed before staining with an anti-FLAG antibody and analyzed by confocal microscopy. The fluorescence corresponding to each protein along the lines traced over condensates is indicated on the graphs. **H)** SINTBAD^KO^ HEK293T cells were co-transfected with plasmids encoding GFP-TBK1 and mC-NAP1 (FL or 210-392) for 24 h. After immunoprecipitation of TBK1 with an anti-GFP antibody, cell lysates (WCL) and immunoprecipitated proteins (IP) were analyzed by WB with anti-TBK1, anti-mC and anti-GAPDH antibodies. On microscopy images, square boxes have been magnified (3X) at the top left. Scale bars: 10µm and 2 µm for insets.

### TBK1 interaction with adaptor proteins is required for its recruitment into NAP1 condensates and disruption of SINTBAD and TANK condensates

We then characterized the role of the interaction between TBK1 and adaptor proteins in condensate formation. For this, we first co-expressed a mutant of TBK1 deleted of its ABD (GFP-TBK1 1-668 – Figure S3A) with the full-length (FL) adaptor proteins fused to mC. The absence of interaction between GFP-TBK1 1-668 and adaptor proteins was verified by co-immunoprecipitation in HEK293T cells (Figure S3B). The analysis of the cellular localization of the proteins revealed that GFP-TBK1 1-668 was not recruited into NAP1 condensates and did not impact the condensation of SINTBAD and TANK (Figure 3C). Of note, GFP-TBK1 1-668 was as effective as GFP-TBK1 in inducing the IFN-β pathway indicating that this deletion mutant was correctly folded and functional (Figure S3C), as previously described (24).

We then performed the reverse experiments by co-expressing GFP-TBK1 with deletion mutants of adaptor proteins (both HD alone and TBD-CTD) fused to mC. For each TBK1- adaptor protein, we verified by co-immunoprecipitation assays that TBK1 binds TBD-CTD but not HD (Figure S3D). The analysis of protein localization revealed that TBK1 had no negative impact on condensates formed through the HD of either SINTBAD or TANK (Figures 3D and 3E). Moreover, TBK1 was not recruited into the condensates formed by the HD of NAP1 (Figure 3F). Surprisingly, co-expression of TBK1 and NAP1 210-392 (corresponding to TBD-CTD) resulted in condensate formation, although neither protein alone was able to form condensates (Figures 1D and 3B). However, these condensates did not form in SINTBAD^KO^ HEK293T cells, suggesting that SINTBAD was required for their formation (Figure 3G). Indeed, co-expression of GFP-SINTBAD with mC-NAP1 210-392 and FLAG-TBK1 led to the co-localization of the three proteins in the same condensates in the parental HEK293T cell line and restored condensates formation in the SINTBAD^KO^ HEK293T (Figure 3G). Of note, the phenotype observed in SINTBAD^KO^ HEK293T cells was not due to a lack of interaction between TBK1 and NAP1 210-392 (Figure 3H).

Taken together, these experiments demonstrated that the interaction between TBK1 and NAP1 was necessary for TBK1 to accumulate into NAP1 condensates. Additionally, the ability of TBK1 to disrupt SINTBAD and TANK condensates required its interaction with the adaptor proteins.

### NAP1 exerts both an early activating and a late limiting effect on the TBK1-dependent IFN-β induction pathway

To assess the potential role of adaptor proteins and their condensates in the induction of the IFN-β pathway, we infected NAP1^KO^, SINTBAD^KO^ or TANK^KO^ cell lines (HEK293T or HAP1) with RABV and quantified the expression of *IFN-β* mRNAs by RT-qPCR 16 hours post-infection (p.i.). We observed a fourfold increased induction of *IFN-β* mRNAs in NAP1^KO^ cell lines compared to parental cell lines, while we did not observe such an increase in SINTBAD^KO^ or TANK^KO^ cell lines (Figures 4A and S4A). Likewise, transfecting NAP1^KO^ cell lines either with a plasmid encoding ΔRIG-I (a constitutively active form of RIG-I) or GFP-TBK1 (leading to the kinase overexpression and consequently to its autophosphorylation) to activate the IFN-β induction pathway, led to a significant increase in *IFN-β* mRNAs compared to parental cell lines (Figures 4B-C and S4B-C). However, no significant effect on IFN-β induction was observed in TANK^KO^ or SINTBAD^KO^ cell lines (Figures 4B-C and S4B-C). Finally, when the IFN-β induction pathway was activated by expressing a phosphomimetic version of IRF3, IRF3-5D (30), no difference in IFN-β induction was observed between the parental and the KO cell lines (Figures 4D and S4D). Taken together, these data (obtained on distinct NAP1^KO^ cell types constructed using distinct RNA guides) indicated that NAP1 limits the induction of IFN-β at a stage upstream of IRF3 phosphorylation.

**Fig. 4.**
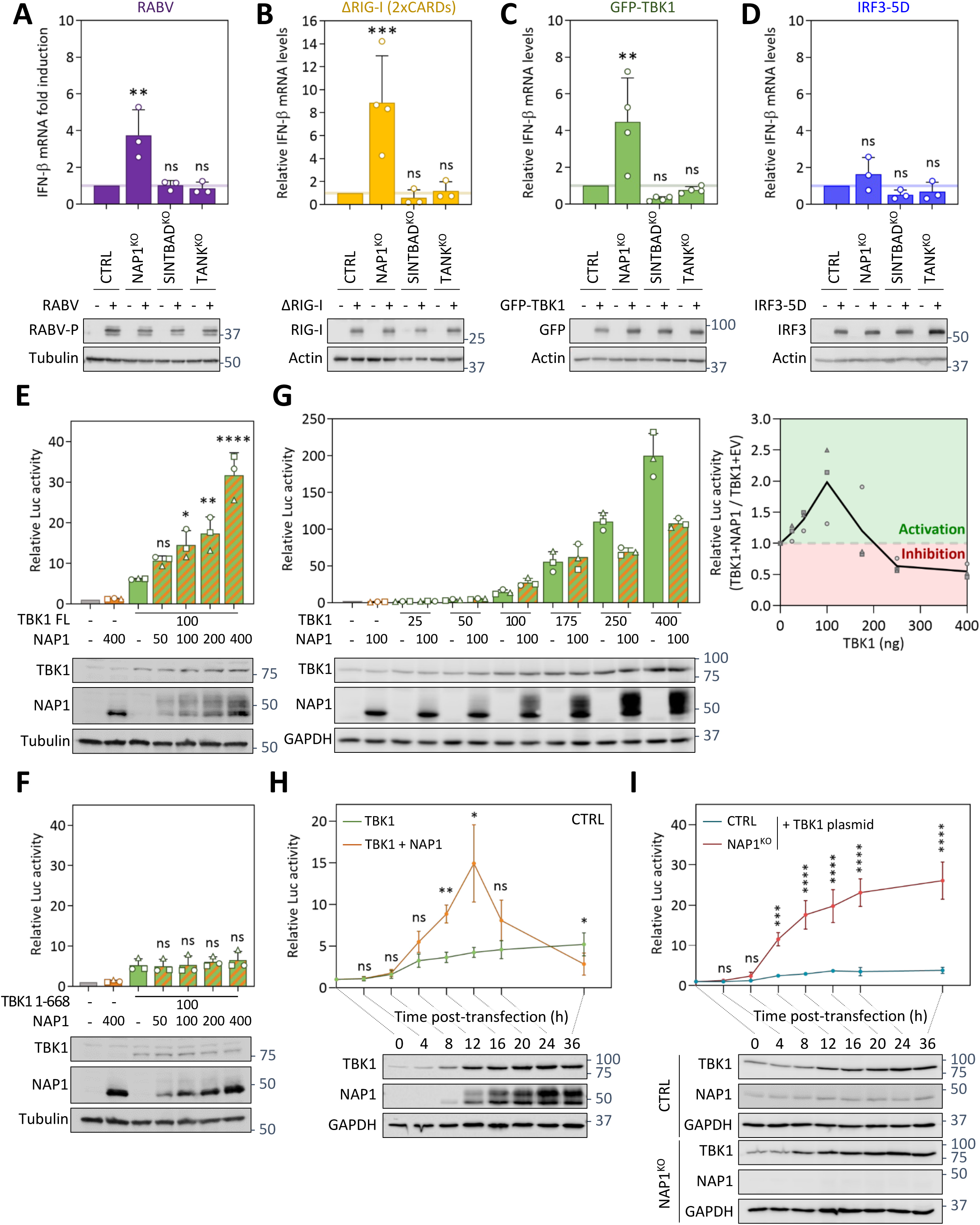
NAP1 has a dual effect on TBK1 activity: inducer at the early stage and limiter at the later stage. **A-D)** Parental (CTRL) and KO HEK293T cells (NAP1^KO^, SINTBAD^KO^, and TANK^KO^) were infected (MOI=3) with RABV (**A**), or transfected either with ΔRIG-I (**B**), GFP-TBK1 (**C**) or IRF3-5D (**D**) plasmids. At 16 h p.i. or 24 h p.t., cells were lysed and the relative levels of *IFN-β* mRNAs to *GAPDH* mRNAs were quantified by RT-qPCR. For each KO cell line, the *IFN-β* mRNA fold induction relative to the control cells is indicated. The bars represent the mean ± SD of at least 3 independent experiments and the corresponding data points are shown. Cell lysates were also analyzed by WB using anti-P, anti-RIG-I, anti-GFP, anti-IRF3, anti-Actin, or anti-Tubulin antibodies. **E-F)** NAP1^KO^ HEK293T cells were co-transfected with a fixed amount (100ng) of plasmids encoding TBK1 FL (**E**) or 1-668 (**F**) and increasing amounts (0-400 ng) of plasmids encoding NAP1 as well as the plasmids of the Luc reporter assay. After 24 h, cells were lysed and the induction of the IFN pathway was assessed using the Luc reporter assay (activation of the *IFN-β* promoter). The IFN-β Luc fold induction relative to the empty vector condition is indicated. The bars represent the mean ± SD of 3 independent experiments and the corresponding data points are shown. Cell lysates were also analyzed by WB using anti-TBK1, anti-NAP1 and anti-Tubulin antibodies. **G)** HEK293T cells were co-transfected with a fixed amount (100ng) of plasmids encoding NAP1 (or EV) and increasing amounts (0-400ng) of plasmids encoding TBK1 as well as the plasmids of the Luc reporter assay. After 24 h, cells were lysed and the induction of the IFN pathway was assessed using the Luc reporter assay (activation of the IFN-β promoter). *Left*, The IFN-β Luc fold induction relative to the empty vector condition is indicated. The bars represent the mean ± SD of 3 independent experiments and the corresponding data points are shown. Cell lysates were also analyzed by WB using anti-TBK1, anti-NAP1, and anti-GAPDH antibodies. *Right*, the dual role of NAP1 on TBK1 activity was shown by calculating the ratio of Luc activities between the condition where TBK1 and NAP1 are co-expressed with the condition where TBK1 is expressed alone. These ratios are calculated for each amount of plasmid encoding TBK1 used. The part of the graph corresponding to activation (resp. inhibition) is in green (resp. pink). **H)** HEK293T cells were transfected with plasmids encoding TBK1 (100 ng – in green) or co-transfected with plasmids encoding TBK1 and NAP1 (100 ng each – in orange) as well as the plasmids of the Luc reporter assay. At each indicated time point, cells were lysed and the induction of the IFN pathway relative to the initial time point (t=0) was assessed using the Luc reporter assay (activation of the IFN-β promoter). Each point on the graph represents the mean ± SD of 3 independent experiments. Cell lysates were also analyzed by WB using anti-TBK1, anti-NAP1, and anti-GAPDH antibodies. **I)** Parental (CTRL - in blue) and NAP1^KO^ (in red) HEK293T cells were transfected with plasmids encoding TBK1 (100 ng) and the plasmids of the Luc reporter assay. At each indicated time point, cells were lysed and the induction of the IFN pathway relative to the initial time point (t=0) was assessed using the Luc reporter assay (activation of the IFN-β promoter). Each point on the graph represents the mean ± SD of three independent experiments. Cell lysates were also analyzed by WB using anti-TBK1, anti-NAP1, and anti-GAPDH antibodies. ns =non-significant; *P < 0.05; **P < 0.01; ***P < 0.001; ****P < 0.0001 (ANOVA [A-D and I] or Student t test [E, F and H]).

This finding was somewhat surprising, as previous luciferase (Luc) reporter gene assays demonstrated that co-expression of NAP1 and TBK1 enhances the activation of the TBK1-dependent IFN-β induction pathway (19). Indeed, using the same Luc assay, when we co-transfected 100 ng of TBK1-encoding plasmid with increasing amounts of NAP1-encoding plasmid (0 to 400 ng), we observed a NAP1-dose dependent enhancement of the activation of the IFN-β induction pathway (Figure 4E). Of note, this enhancement required the interaction between NAP1 and TBK1 as it was not observed when the TBK1 1-668 mutant (which does not bind NAP1 – Figure S3D) was expressed instead of the TBK1 FL (Figure 4F).

We also carried out the opposite experiment in which we co-transfected 100 ng of NAP1-encoding plasmids with increasing amounts of TBK1-encoding plasmids (0 to 400 ng). We observed that up to 175 ng of TBK1 plasmids, NAP1 enhanced the TBK1-induced activation of the IFN-β induction pathway while above 250 ng of TBK1 plasmids, NAP1 limited the IFN-β response (Figure 4G).

Overall, these experiments were consistent with a potential dual role of NAP1. On the one hand, it contributes to increasing TBK1 activity; on the other hand, it can limit the induction of the IFN-β pathway. In the latter experiment (Figure 4G), at a high TBK1 concentration, the kinase spontaneously homodimerizes which promotes its activation, by intermolecular transphosphorylation (12,31,32). Under these conditions, NAP1 probably only marginally activates the kinase, highlighting its inhibitory effect on IFN-β induction.

We hypothesized that the activating effect of NAP1 would manifest early in the pathway induction and that the limiting effect would emerge later when it is necessary to stop the IFN-β pathway, and thus avoid excessive IFN-β production. To validate this hypothesis, we performed a Luc assay in NAP1^KO^ HEK293T cells, in which we compared the kinetics of IFN-β induction after transfection with a plasmid encoding TBK1 and after co-transfection with plasmids encoding TBK1 and NAP1 (Figure 4H). Our results showed that co-expression of NAP1 and TBK1 significantly enhanced IFN-β induction up to 20 h post-transfection (p.t.). However, after 20 h, the enhancement was less pronounced and even, after 36 h, co-expression of NAP1 had a negative effect on IFN-β induction (Figure 4H). This confirmed that over time, NAP1 was shifting from an activator of the IFN-β induction pathway toward a limiter of this pathway.

Finally, we compared the kinetics of IFN-β induction between the NAP1^KO^ HEK293T cell line and the parental cell line, both transfected with a plasmid encoding TBK1 (Figure 4I). We observed a progressive increase of IFN induction over time in NAP1^KO^ cells, while this induction was very rapidly limited in parental cells that expressed NAP1, highlighting the limiting role of endogenous NAP1 in the process (Figure 4I).

### Concentration of TBK1 into NAP1 condensates limits the induction of the IFN-β pathway

Activated TBK1 phosphorylates transcription factors IRF3 and/or IRF7, resulting in their homo- or heterodimerization and subsequent nuclear translocation, thereby inducing type I IFN. To investigate the impact of NAP1 on this process, we transfected cells with a plasmid expressing either mC-NAP1 or GFP-TBK1, or with both plasmids leading to co-expression of the two proteins. We then investigated the localization of endogenous IRF3 by IF in the transfected cells. The ectopic expression of GFP-TBK1 (FL or 1-668) alone induced translocation and accumulation of IRF3 into the nucleus, a sign of its activation, while the ectopic expression of mC-NAP1 (or its deletion mutants) had no effect (Figures 5A-B). When both GFP-TBK1 and mC-NAP1 were ectopically expressed, we observed GFP-TBK1 concentration in mC-NAP1 condensates accompanied by a cytoplasmic localization of IRF3, again demonstrating the inhibitory effect of NAP1 on IFN induction (Figures 5A-B).

**Fig. 5.**
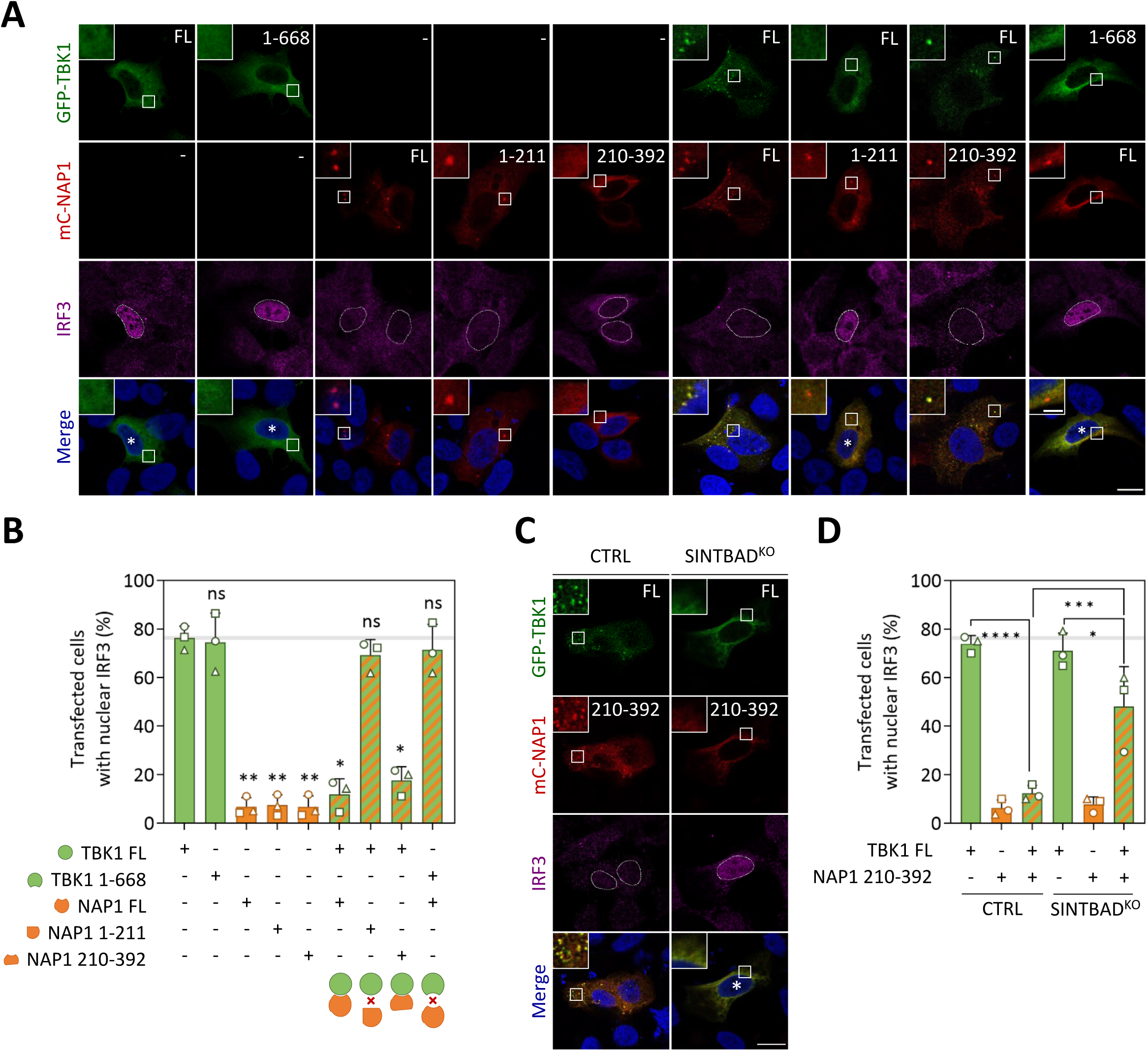
TBK1 accumulation inside NAP1 condensates inhibits IRF3 activation. **A-B)** HAP1 cells were single- or co-transfected with plasmids encoding GFP-TBK1 (FL or 1-668) and mC-NAP1 (FL, 1-211 or 210-392). After 24 h, cells were fixed before staining with an anti-IRF3 antibody. (**A)** Analysis by confocal microscopy. (**B)** Percentage of transfected cells with IRF3 nuclear localization is indicated. The bars represent the mean ± SD and data points represent replicates experiments (n=3). The red crosses on the drawings below the diagram indicate the absence of interaction between the indicated proteins. **C-D)** Parental (CTRL) and SINTBAD^KO^ HAP1 cells were single and co-transfected with plasmids encoding GFP-TBK1 FL and mC-NAP1 210-392. After 24 h, cells were fixed before staining with an anti-IRF3 antibody. (**C)** Analysis by confocal microscopy. (**D)** Percentage of transfected cells with IRF3 nuclear localization is shown. The bars represent the mean ± SD and data points represent replicates experiments (n=3). On microscopy images, square boxes have been magnified (3X) at the top left. Dotted lines delimit the nucleus and asterisks indicate cells exhibiting an IRF3 nuclear localization. Scale bars: 10 µm and 2 µm for insets. ns =non-significant; *P < 0.05 ; **P < 0.01 ; ***P < 0.001 ; ****P < 0.0001 (ANOVA test).

To elucidate the mechanism by which NAP1 inhibits IFN induction, we co-transfected HAP1 cells to ectopically express TBK1 and NAP1 1-211 (corresponding to HD), TBK1 and NAP1 210-392 (corresponding to TBD-CTD), and TBK1 1-668 and NAP1. We found that inhibition of IRF3 nuclear translocation only occurred when NAP1 (or its deletion mutants) formed condensates concentrating TBK1 (Figures 5A-B). This suggested that the concentration of TBK1 into NAP1 condensates was crucial. Nevertheless, as we could not dissociate TBK1 and NAP1 interaction from TBK1 accumulation into NAP1 condensates, we could not formally exclude that the inhibitory effect of NAP1 was solely due to its interaction with the kinase.

To address this point, we took advantage of our observation that the co-expression of TBK1 and the NAP1 210-392 deletion mutant resulted in the formation of condensates in parental but not in SINTBAD^KO^ HEK293T cells (Figures 3G). Remarkably, inhibition of IRF3 nuclear translocation by mC-NAP1 210-392 was only observed in the parental HEK293T cell line in which condensates are formed (Figures 5C-D). This definitively demonstrated that the interaction between TBK1 and NAP1 210-392 is insufficient for NAP1 210-392 to exert its inhibitory effect and that accumulation of TBK1 into condensates is required.

### TBK1 phosphorylates NAP1 on several sites

NAP1 is known to be phosphorylated on several sites by TBK1 (33–35). Consistently, when lysates from HEK293T cells, in which mC-NAP1 was expressed along with GFP-TBK1, were analyzed by Western Blot (WB), mC-NAP1 was detected as multiple bands migrating slower than the one observed when mc-NAP1 was expressed alone (Figure 6A). These bands were not observed when NAP1 was co-expressed with enzymatically inactive TBK1 mutants - K38A which is catalytically inactive (36), S172A which cannot be phosphorylated at position 172, a key autophosphorylation event required to activate the kinase (36), and Y179A, because phosphorylation of tyrosine 179 is required for subsequent autophosphorylation of TBK1 at serine 172 (37) - (Figure 6A). Of note, the interaction between the two proteins involving the C-terminal part of TBK1 was not required for NAP1 phosphorylation, as co-expression of mC-NAP1 and GFP-TBK1 1-668 resulted in the same migration profile (Figure 6A).

**Fig. 6.**
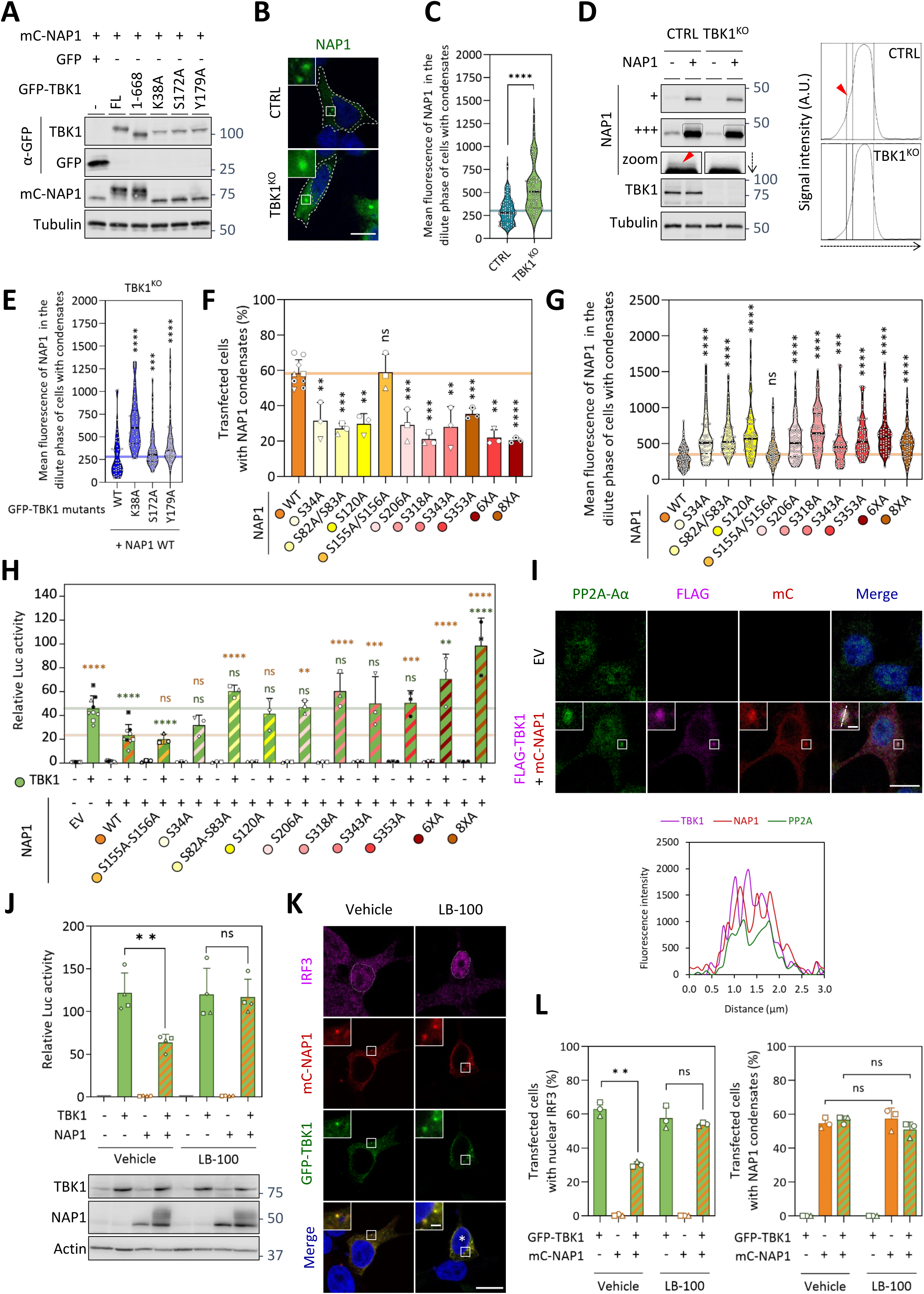
TBK1-dependent phosphorylation of NAP1 induces the formation of condensates that concentrate the kinase and the PP2A phosphatase complex. **A)** HEK293T cells were co-transfected with plasmids encoding mC-NAP1 and GFP-TBK1 WT and mutants (1-668, K38A, S172A or Y179A) or GFP. After 24 h, cells were lysed and analyzed by WB using anti-GFP, anti-mC and anti-Tubulin antibodies. **B-C)** Parental (CTRL) and TBK1^KO^ HEK293T cells were transfected with plasmids encoding NAP1. After 24 h, cells were fixed before staining with an anti-NAP1 antibody. **(B)** Analysis by confocal microscopy. **(C)** Quantification of the mean fluorescence of NAP1 in the dilute phase of transfected cells containing condensates. The violin plots show the median and quartiles from three independent experiments. Each point represents the value for one cell and at least 20 cells per experiment (n=3) were quantified. **D)** Parental (CTRL) and TBK1^KO^ HEK293T cells were transfected with plasmids encoding NAP1 (or EV). *Left,* after 24 h, cells were lysed and analyzed by WB using anti-TBK1, anti-NAP1 and anti-Tubulin antibodies. Bands corresponding to NAP1 are shown with two exposure times (+ and +++) and have been enlarged (zoom) to visualize the phosphorylated form (red arrowhead). *Right*, using ImageJ software, the signal intensity (in arbitrary unit – A.U.) of NAP1’s band in each cell line have been measured along the direction of the dotted arrow. The plots of signal intensity along the dotted arrow are presented. The shoulder corresponding to the phosphorylated form is indicated (red arrowhead). **E)** TBK1^KO^ HEK293T cells were co-transfected with plasmids encoding NAP1 and GFP-TBK1 WT or mutants (K38A, S172A or Y179A). After 24 h, cells were fixed before staining with an anti-NAP1 antibody. Quantification of the mean fluorescence of NAP1 in the dilute phase of co-transfected cells containing condensates. The violin plots show the median and quartiles from 3 independent experiments. Each point represents the value for one cell and at least 20 cells per experiment were quantified. Representative confocal images were presented in Figure S6A. **F-G)** NAP1^KO^ HEK293T cells were transfected with plasmids encoding indicated NAP1 phosphorylation mutants. After 24 h, cells were fixed before staining with an anti-NAP1 antibody. **(G)** Percentage of transfected cells with NAP1 condensates. The bars represent the mean ± SD and data points represent replicates experiments (n=3). **(H)** Quantification of the mean fluorescence of NAP1 in the dilute phase of transfected cells containing condensates. The violin plots show the median and quartiles from 3 independent experiments. Each point represents the value for one cell and at least 20 cells per experiment were quantified. Representative confocal images were presented in Figure S6C. **H)** NAP1^KO^ HEK293T cells were co-transfected with plasmids encoding TBK1 (or EV) and indicated NAP1 phosphorylation mutants as well as the plasmids of the Luc reporter assay. After 24 h, cells were lysed and the induction of the IFN pathway was assessed using the Luc reporter assay (activation of the IFN-β promoter). The fold induction relative to EV-transfected cells condition is indicated. The bars represent the mean ± SD of 3 independent experiments. The statistical significances of the difference with the condition TBK1 + EV (resp. TBK1 + NAP1 WT) are indicated in green (resp. in orange). **I)** HEK293T cells were co-transfected or not (EV) with plasmids encoding FLAG-TBK1 and mC-NAP1. After 24 h, cells were fixed, stained with anti-PP2A-Aα and anti-FLAG antibodies and analyzed by confocal microscopy. The fluorescence corresponding to each protein along the line traced over condensates is indicated on the graph. **J)** NAP1^KO^ HEK293T cells were co-transfected with plasmids encoding TBK1 (or EV) and NAP1 (or EV) as well as the plasmids of the Luc reporter assay. After 20 h, cells were treated or not (Vehicle) with LB-100 for 4 additional hours. Then, cells were lysed and the induction of the IFN pathway was assessed using the Luc reporter assay (activation of the IFN-β promoter). The fold induction relative to EV-transfected cells conditions exposed to the vehicle is indicated. The bars represent the mean ± SD of four independent experiments. Cell lysates were also analyzed by WB using anti-TBK1, anti-NAP1 and anti-Actin antibodies. **K-L)** HEK293T cells were co-transfected with plasmids encoding GFP-TBK1 and mC-NAP1. After 20 h, cells were treated or not (Vehicle) with LB-100 for 4 additional hours. Then, cells were fixed before staining with an anti-IRF3 antibody. **K)** Analysis by confocal microscopy. **L)** *Left*, percentage of transfected cells with IRF3 nuclear localization. *Right*, percentage of transfected cells with NAP1 condensates. The bars represent the mean ± SD and data points represent replicates experiments (n=3). On microscopy images, square boxes have been magnified (3X) at the top left. Yellow dotted lines delimit the nucleus, while white dotted lines delimit the cytoplasm. Scale bars: 10µm and 2 µm for insets. ns =non-significant; *P < 0.05; **P < 0.01; ***P < 0.001; ***P < 0.0001 (Student t-test [**B**, **J** and **L**], Kolmogorov-Smirnov test [**C**, **E** and **G**] or ANOVA [**F** and **H**]).

To identify the phosphorylation sites on NAP1, we immunoprecipitated GFP-NAP1 from HEK293T cells co-expressing GFP-NAP1 and mC-TBK1. Immunoprecipitates were analyzed by SDS-PAGE (Figure S5A) and WB (Figure S5B). Bands corresponding to NAP1 were excised and digested with trypsin protease before analysis by LC-MS/MS. This approach identified serine residues 82 (and/or 83), 120, 206, 318 and 353 as phosphorylation sites (Figures S5C-D). It should be noted that residues 82 (and/or 83), 120 and 318 were previously identified as phosphorylation sites, as well as serine residues 34 and 343, which were not detected in our analysis (34).

### NAP1 phosphorylation enhances its ability to form condensates

Several of our observations suggested that TBK1-dependent phosphorylation of NAP1 favors its condensation. First, in cells not exposed to a danger signal, in which TBK1 was not activated, endogenous NAP1 does not form condensates (Figures 1A-B, S1A, and S1C). Second, upon cell transfection with 5’ppp-dsRNA, significantly fewer TBK1^KO^ HEK293T cells contained endogenous NAP1 condensates compared to parental cells (Figure 1B).

We hypothesized that NAP1 phosphorylation lowers its critical concentration required for LLPS. To test this, we transfected parental and TBK1^KO^ HEK293T cells by a plasmid encoding NAP1 and detected the adaptor protein by IF using an anti-NAP1 antibody (Figure 6B). Then, in cells containing condensates, we measured the concentration of NAP1 in the dilute phase (*i.e.* outside the condensates) by assuming that the local concentration of the protein was proportional to the local fluorescence intensity per pixel. This indicated that the concentration of NAP1 in the dilute phase was higher in the TBK1^KO^ HEK293T than in the parental cells (Figure 6C). Indeed, in parental HEK293T cells (but not in TBK1^KO^ HEK293T cells) ectopically expressing NAP1, it was possible to detect a phosphorylated form of NAP1 by WB, although in low quantity (Figure 6D).

To confirm that NAP1 phosphorylation led to a decrease of its critical concentration required to form condensates, we co-transfected TBK1^KO^ HEK293T cells with plasmids encoding NAP1 and either GFP-TBK1 wild type (WT) or kinase inactive mutants (K38A, S172A, Y179A) (Figures 6F and S6A). As expected, these mutants did not induce transcription of *IFN-β* genes (Figure S6B). We observed that the expression of GFP-TBK1 K38A, and to a lesser extent S172A and Y179A, increased the concentration of NAP1 in the dilute phase compared to GFP-TBK1 WT (Figures 6E and S6A). Therefore, the TBK1-dependent phosphorylation of NAP1 decreases NAP1’s critical concentration to form condensates.

To further investigate the role of NAP1 phosphorylation, we constructed plasmids encoding NAP1 mutants, in which phosphorylable serine residues were mutated to alanine, either individually or in combination (Figure S5C). NAP1^KO^ HEK293T cells were transfected with these plasmids before IF analysis using an anti-NAP1 antibody (Figures 6F-G and S6C). For each NAP1 mutant, we assessed the proportion of cells containing condensates (Figure 6F), and when condensates were present, we also measured NAP1 concentration in the dilute phase (Figure 6G). All NAP1 phosphorylation mutants showed a reduced ability to induce LLPS and higher concentration in the dilute phase compared to NAP1 WT. The fact that all mutants negatively impacted the ability of NAP1 to form condensates was somewhat puzzling. We, therefore, constructed the NAP1 double mutant (S155A/S156A) because serine residues 155 and 156 had not been identified as phosphorylated. This mutant had a phenotype similar to NAP1 WT (Figure 6F-G and S6C). There is, therefore, a correlation between NAP1 phosphorylation at a global level (not involving a specific site) and its ability to form condensates.

Finally, we assessed the impact of NAP1 phosphorylation mutants on IFN-β induction using the Luc assay in NAP1^KO^ HEK293T cells. These cells were co-transfected with plasmids encoding TBK1 (to activate the IFN-β induction pathway), and either NAP1 WT, or NAP1 phosphorylation mutants (Figure 6H). As expected, the S155A/S156A mutant behaved like NAP1 WT and inhibited TBK1 activity. All other NAP1 single and double mutants having a defect in condensate formation had a significantly lower ability to inhibit TBK1 activity than NAP1 WT. Remarkably, in the case of mutants 6XA and 8XA, only the activating effect of the mutants on the IFN induction pathway was observed. Taken together, these data indicate that NAP1 phosphorylation is required for its inhibitory effect but has no impact on its activating effect.

Overall, these results demonstrate that TBK1-dependent phosphorylation of NAP1 enhances its ability to form condensates, which in turn downregulates the TBK1-dependent IFN-β pathway. This is also consistent with the data shown in Figure 4G, where the inhibitory effect of NAP1 was detected when NAP1 phosphorylation was observed.

### PP2A phosphatase complex is concentrated in NAP1 condensates to deactivate TBK1

Several of our experiments suggested that the limiting effect of NAP1 on IFN-β induction was not due to a stoichiometric ratio between NAP1 and TBK1, as this effect was particularly observed at high concentrations of TBK1 (Figures 4G). Furthermore, we showed by FRAP experiments that TBK1 can shuttle between the cytoplasm and NAP1 condensates with a mobile fraction of ∼60% and rapid recovery of the fluorescence signal (t_1/2_∼2s) (Figure S6D). Such a short residence time suggested that TBK1 sequestration alone could not fully explain the inhibitory action of NAP1, and that the condensates might contain a phosphatase that could dephosphorylate TBK1.

A good candidate for this role could be a member of the PP2A family, one of the phosphatase families known to dephosphorylate TBK1 (38). Indeed, endogenous PP2A-Aα, the scaffold subunit of the PP2A family, co-localized with condensates formed by ectopically expressed FLAG-TBK1 and mC-NAP1 (Figure 6I). Endogenous PP2A-Aα condensates were also observed when cells were exposed to danger signals such as RABV infection (Figure S6E) or transfection with 5’ppp-dsRNA (Figure S6F). Due to antibody limitations, we were unable to assess the localization of endogenous TBK1 and NAP1 in these experiments. By co-IP experiment, we also investigated a potential interaction between the phosphatase and TBK1 or NAP1. As we did not have a suitable antibody directed against PP2A for this purpose, we constructed a GFP-tagged version of the catalytic subunit of the phosphatase family (GFP-PP2A-Cα). We then immunoprecipitated the latter and found that NAP1, but not TBK1, was associated with GFP-PP2A-Cα (Figure S6G). Indeed, in live imaging, we observed that GFP-PP2A-Cα accumulated inside NAP1 condensates, regardless of the presence of TBK1 (Figure S6H).

To demonstrate the role of the PP2A family in the inhibition of the TBK1-dependent IFN-β induction pathway, cells were treated with LB-100, a potent and specific PP2A inhibitor (39). This abolished the inhibitory effect of NAP1 on the TBK1-dependent induction of the IFN-β promoter (Figure 6J) and prevented the activation of IRF3 (Figures 6K-L). Importantly, this effect is not due to a disruption of NAP1 condensates following LB-100 treatment, as we still observed the same proportion of cells containing NAP1 condensates (Figure 6L right panel).

Overall, our results demonstrated that NAP1 condensates concentrate both PP2A and TBK1, allowing PP2A to dephosphorylate TBK1, leading to its deactivation and limiting the production of IFN-β.

### NAP1 variants are found in patients with immune-related dysregulations

We searched for rare NAP1 variants in an in-house exome dataset generated from a cohort of > 2000 patients presenting with early-onset autoimmune diseases, including SLE (>200 patients) and idiopathic interferonopathy (>20 patients). We found *AZI2* mono-allelic private deleterious mutations in three patients (2 SLE and 1 idiopathic interferonopathy). Two mutations (V84M and K109E) were located in the HD domain and one (P302H) was located in an IDD (Figure 7A). All variants exhibit a high deleterious predicting score, supporting a potential causal role in the pathology (Figure S7A). Patient A (p.V84M) presented with a severe idiopathic early-onset interferonopathy-like disease with elevated type I IFN plasmatic concentration, brain calcification, interstitial lung disease, and lymphoproliferation with auto-immune haemolytic anemia (Supplementary information). Patient B (p.K109E) developed early-onset (<5 years of age) SLE and patient C (p.P302H) was affected by a syndromic SLE (Supplementary information). All known genes involved in monogenic interferonopathy were screened and no other mutation could be found. Therefore, these mutations in NAP1 could define a new monogenic predisposition to interferon-related diseases (interferonopathy and SLE) in agreement with our data indicating a role for NAP1 in the negative regulation of the IFN induction pathway.

**Fig. 7.**
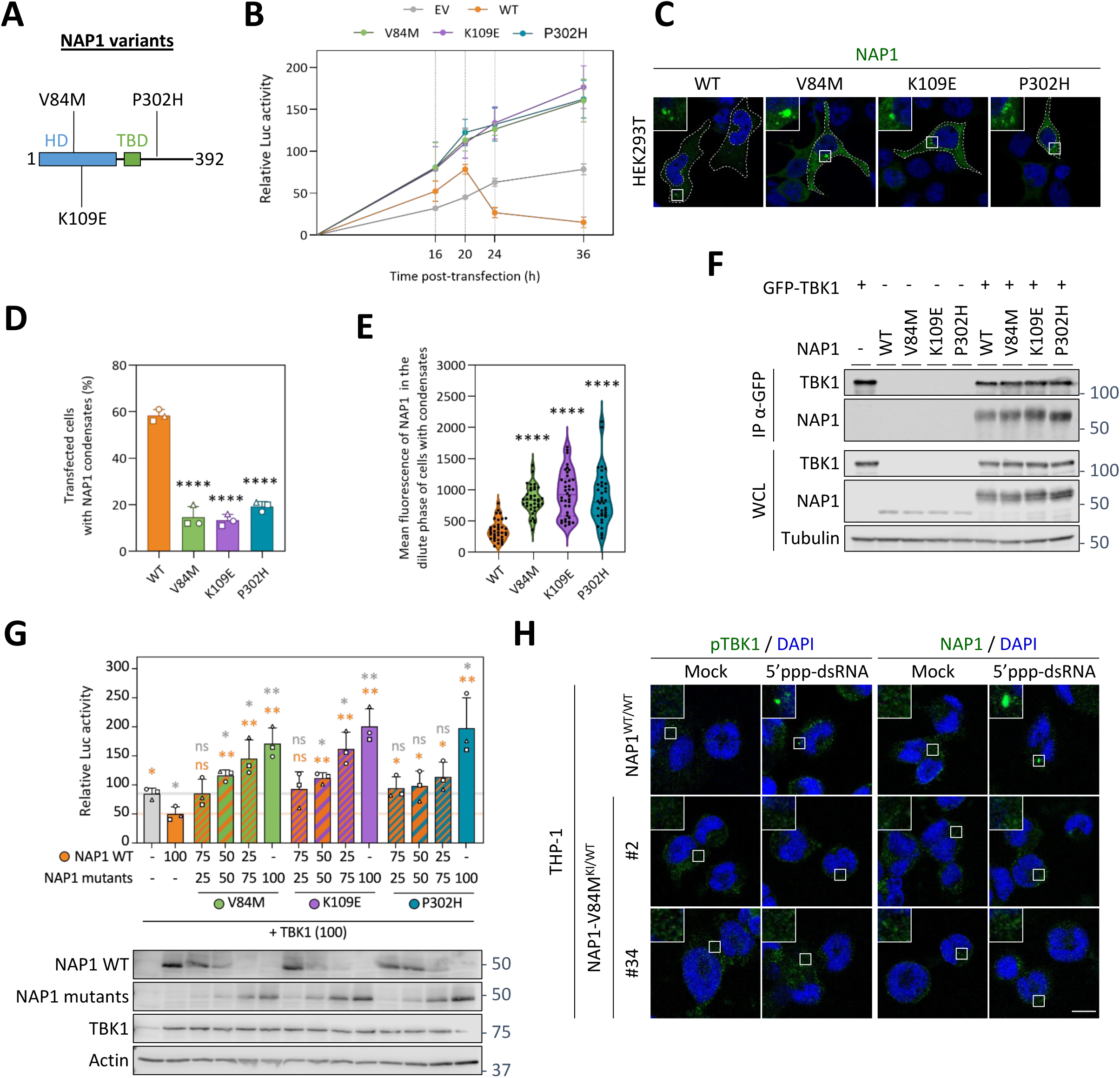
NAP1 variants, associated with immune diseases in patients, have a defect in condensate formation and the downregulation of TBK1-dependent IFN-β pathway. **A)** NAP1 variants identified in immune diseases. **B)** NAP1^KO^ HEK293T cells were co-transfected with plasmids encoding TBK1 (100 ng) and NAP1 mutants (100 ng each – or EV) as well as the plasmids of the Luc reporter assay. At each indicated time point, cells were lysed and the induction of the IFN pathway was assessed using the Luc reporter assay (activation of the IFN-β promoter). Each point on the graph represents the mean ± SD of three independent experiments. **C-E)** NAP1^KO^ HEK293T cells were transfected with plasmids encoding NAP1 variants (WT, V84M, K109E and P302H). After 24 h, cells were fixed and immunostained with an anti-NAP1 antibody. **(C)** Analysis by confocal microscopy. **(D)** Percentage of transfected cells with NAP1 condensates. The bars represent the mean ± SD and data points represent replicates experiments (n=3). **(E)** Quantification of the mean fluorescence of NAP1 in the dilute phase of transfected cells containing condensates. The violin plots show the median and quartiles from 3 independent experiments. Each point represents the value for one cell and at least 20 cells per experiment were quantified. **F)** HEK293T cells were co-transfected with plasmids encoding indicated NAP1 mutants and GFP-TBK1 (or EV) for 24 h. After immunoprecipitation of TBK1 with an anti-GFP antibody, cell lysates (WCL) and immunoprecipitated proteins (IP) were analyzed by WB with anti-TBK1, anti-NAP1 and anti-Tubulin antibodies. **G)** NAP1^KO^ HEK293T cells were co-transfected with plasmids encoding TBK1 (100 ng) and with the different ratio of plasmids encoding FLAG-NAP1 WT or MYC-NAP1 variants (total of 100 ng) as well as the plasmids of the Luc reporter assay. After 24 h, cells were lysed and the induction of the IFN pathway was assessed using the Luc reporter assay (activation of the IFN-β promoter). The fold induction relative to the EV condition is indicated. The bars represent the mean ± SD of 3 independent experiments. The statistical significances of the difference with the condition TBK1 alone (resp. TBK1 + NAP1 WT) are indicated in grey (resp. in orange). Cell lysates were also analyzed by WB using anti-FLAG, anti-MYC, anti-TBK1 and anti-Actin antibodies. **H)** THP-1 NAP1^WT/WT^ and THP-1 NAP1 V84M^KI/WT^ (clones #2 et #34) cells were incubated with medium containing LyoVec alone (Mock) or transfected with 5’ppp-dsRNA using LyoVec (5’ppp-dsRNA). After 6 h, cells were fixed before staining with anti-pTBK1 (S172) or anti-NAP1 antibody and analyzed by confocal microscopy. On microscopy images, square boxes have been magnified (3X) at the top left. Yellow dotted lines delimit the nucleus, while white dotted lines delimit the cytoplasm. Numbers at the top right indicate the rate of transfected cells with NAP1 condensates. Scale bars: 10µm and 2 µm for insets. ns =non-significant; *P < 0.05; **P < 0.01, ***P < 0.001 (ANOVA [**D**, **E** and **H**] or Kolmogorov-Smirnov test [**F**]).

To further support the causal role of these variants, we constructed pcDNA plasmids encoding these NAP1 variants and assessed them in our cellular models. We used the Luc assay to characterize the impact of the expression of NAP1 mutants V84M, K109E and P302H in NAP1^KO^ HEK293T cells on the IFN-β induction pathway activated by TBK1 ectopic expression. At early time points (*i.e.* < 20 h p.t.), all NAP1 variants induced more efficiently the IFN-β pathway than NAP1 WT and, more importantly, they were unable to restrict this pathway at later time points (*i.e.* > 24 h p.t. - Figure 7B).

Corroborating these observations, the proportion of cells with NAP1-condensates was lower in cells expressing NAP1 variants (Figures 7C-D). Furthermore, when condensates were observed, the concentration of NAP1 variants in the dilute phase (evaluated by the mean fluorescence intensity) was significantly higher (almost 2.5 times more) for NAP1 variants V84M, K109E and P302H than for NAP1 WT (Figure 7E). As NAP1 variants still interacted with TBK1 (Figure 7F), the increase in their critical concentration was most likely the cause of the observed phenotype.

NAP1 variants identified in patients resulted from mono-allelic mutations, meaning that the patients were heterozygous, and suggesting that the mutations could be dominant. To confirm this, we conducted the same Luc assay as described in Figure 7B, where we transfected different ratios of NAP1 variants *vs* the WT (Figure 7G). Cells were lysed at 24 h p.t. to be in the condition where NAP1 WT acts as an inhibitor of TBK1 activity (Figure 7B). For each NAP1 variant, we observed that the lowest ratio of plasmid encoding NAP1 variants (25 ng *vs* 75 ng of NAP1 WT) resulted in a loss of NAP1 WT inhibitory activity (Figure 7G). As we increased the proportion of plasmids encoding NAP1 variants, we noticed an increase in TBK1-dependent induction of the IFN-β pathway. This result demonstrated the dominant effect of mutations V84M, K109E, and P302H of NAP1.

To evaluate the role of endogenous expression of the NAP1 V84M variant, we introduced the mutation in THP-1 and CAL-1 cell lines using a CRISPR/Cas9 editing approach. We obtained heterozygous THP-1 NAP1-V84M^KI/WT^ and homozygous CAL-1 NAP1-V84M^KI/KI^ cell lines (Figures S7B and S7C). We transfected the control and knock-in (KI) cell lines with 5’ppp-dsRNA and assessed the formation of TBK1 and NAP1 condensates in PMA-differentiated THP-1 cells (Figure 7H) and CAL-1 cells (Figure S7D) by IF. TBK1 and NAP1 condensates were only observed in homozygous cells expressing a WT version of NAP1, confirming that the mutation V84M abrogates condensate formation and has a dominant phenotype.

Taken together, the data showed that these mutations in the *AZI2* gene affect NAP1’s ability to form condensates, a characteristic that likely contributes to the immune diseases described in these patients.

## Discussion

The formation of cellular condensates, whether formed by LLPS or other mechanisms, is essential for the regulation of several physiological processes(40,41). It now appears that condensates play a role throughout the IFN-induction pathway. As examples, it has been shown that cGAS forms liquid organelles with DNA (42,43), that STING polymerizes upon activation (44) and also makes condensates with stacked ER membrane (45), and finally that RIG-I and MDA-5 are resident proteins of antiviral stress granules(46–49). Moreover, others and we also have recently demonstrated that TBK1 adaptor proteins form condensates upon cell exposure of cells to danger signals (28,50).

Here, complementing these previous observations, we showed that upon activation of the MAVS- or the cGAS-STING-dependent IFN induction pathway, NAP1, SINTBAD and TANK formed condensates exhibiting the hallmarks of transient organelles formed by LLPS (ability to fuse, sensitivity to hypotonic shock and 1,6-HD, rapid recovery of fluorescence after photobleaching). We also identified the amino terminal domain of the adaptor proteins, responsible for their homodimerization, as the driver of the phase separation. Furthermore, we show that NAP1 condensates are able to accumulate TBK1 while the kinase has a negative impact on the formation of condensates formed by TANK and SINTBAD. Both phenomena involve a physical interaction between the kinase and its adaptors.

The major result of this article is the demonstration that the formation of NAP1 condensates limits the IFN induction pathway. More precisely, our data reveal a dual role of NAP1 (Figure S8). First, shortly after the detection of the danger signal, NAP1, by binding to TBK1, increases the activity of the kinase. This is consistent with previous data, which we confirmed here, showing that NAP1 ectopic expression markedly enhances TBK1-mediated IFN-β promoter activation (19). However, since NAP1 is a substrate of the kinase, it is progressively phosphorylated. At some point, the concentration of phosphorylated NAP1 within the cell reaches the critical concentration for condensate formation. These condensates then accumulate TBK1, as well as the phosphatase PP2A (which also binds NAP1), which dephosphorylates the kinase, returning it to its inactive form. This stops the IFN induction pathway.

Interestingly, our systematic mutagenesis of NAP1 phosphorylation sites indicates that the decrease in the critical concentration of NAP1 when it is phosphorylated does not involve a specific site. This suggests that LLPS is instead triggered by the overall charge of certain protein domains in agreement with what is observed for several other proteins capable of inducing phase separations (51,52).

The identification of mono-allelic *AZI2* mutations in patients with autoimmune or inflammatory disorders, such as SLE and idiopathic interferonopathy, suggested a novel monogenic predisposition to IFN-related diseases. The V84M, K109E, and P302H mutations impair the ability of NAP1 to limit the IFN-β induction in an ectopic expression model. Functional assays demonstrated that these variants enhance IFN-β induction at early stages and fail to restrict it later, leading to prolonged immune activation. A key mechanistic insight is the apparent disruption of the ability of NAP1 mutants to form condensates, a feature that we have demonstrated to be essential for modulating TBK1 activity in both an ectopic expression model and in a more physiological knock-in model. Mutant proteins exhibited reduced condensate formation and higher critical concentration while still interacting with TBK1. This alteration likely contributes to excessive IFN-β signaling. Moreover, heterozygous expression of these variants was sufficient to counteract the inhibitory function of NAP1 WT, suggesting a dominant negative effect that renders NAP1 WT unable to restrict IFN activity, likely due to the formation of inactive heterodimers between NAP1 WT and NAP1 V84M. These findings underscore the critical role of NAP1 condensates in immune regulation and suggest that *AZI2* mutations could act as a genetic predisposition to immune dysregulation and related diseases.

This study therefore highlights a new form of regulation involving condensates and shows how a protein can switch from an activating to a limiting state by being the substrate of the kinase it activates. This is possible because the phosphorylated protein has a lower critical concentration than the unmodified protein. In fact, from the moment the IFN induction cascade is triggered, at each instant, the number of NAP1 phosphorylation events is proportional to the amount of activated TBK1 in the cell. NAP1, therefore, behaves as a molecular integrator of total TBK1 activity in the cell since the triggering of the cascade. When a certain threshold is reached, this provokes the formation of condensates in which the kinase is deactivated (Figure S8).

This mode of regulation is probably not specific to the regulation of TBK1 activity by NAP1. Indeed, it has been observed that mutations in the tumor suppressor neurofibromin (NF2/Merlin/schwannomin), a positive regulator of IFN induction, convert the protein into a potent suppressor of cGAS-STING signaling (38). In fact, IRF3, when activated after nucleic acid sensing, induces the formation of condensates containing NF2 mutant, TBK1 and IRF3, as well as the PP2A complex, to dephosphorylate TBK1. In this case, NF2 mutations, by decreasing the critical NF2 concentration required for condensate formation, likely affect a regulatory function similar to that of NAP1. This dysregulation results in tumorigenesis in several organs. Beyond innate immunity, such regulation is probably also exploited in other signaling pathways when it is necessary to limit the duration of the response to a stimulus.

### Limits of the study

In this study, we only identified the role of NAP1 condensates but did not identify any function associated with those formed by SINTBAD and TANK. Since these adaptor proteins also undergo LLPS and interact with TBK1, their potential contribution to the regulation of the IFN pathway or any other pathway in which TBK1 is involved remains an open question.

Since the *AZI2* mutations are inherited from an asymptomatic parent, this indicates that additional modifiers, either intrinsic (genetic or epigenetic) or extrinsic (environmental) are probably involved in the pathophysiological mechanisms underlying disease onset. Further studies, pending the availability of patients’ material, are needed to explore in detail the consequences of these mutations in patient cells.

## Material and methods

### Cell lines

HEK293T cells (human embryonic kidney, ATCC #CRL-3216) were purchased from the ATCC organization (http://www.lgcstandards-atcc.org). CRISPR control (CTRL), TBK1^KO^, NAP1 ^KO^, SINTBAD ^KO,^ and TANK^KO^-HEK293T cells were generated for this study (see section below). All these cells were cultured at 37°C, 5% CO_2_, in Dulbecco’s modified eagle medium (DMEM) (Gibco-Life Technologies) supplemented with 10% fetal calf serum (FCS) (Gibco-Life Technologies), 100 units/mL of Penicillin and 100 mg/mL of Streptomycin (Gibco-Life Technologies). Parental control (CTRL, #C631), TBK1^KO^ (#HZGHC000031c010), NAP1^KO^ (#HZGHC003895c005), SINTBAD^KO^ (HZGHC003903c004), TANK^KO^ (#HZGHC003904c010) and MAVS^KO^ (#HZGHC001456c011) HAP1 cells were purchased from Horizon Discovery (https://horizondiscovery.com/en/engineered-cell-lines). All these cells were cultured at 37°C, 5% CO_2_, in Iscove’s Modified DMEM (IMDM, Corning) supplemented with 10% FCS, 100 units/mL of penicillin, and 100 mg/mL of Streptomycin (AB 1X).

THP-1 cells (ATCC, #TIB-202) were a gift from Yannick Crow laboratory and were initially purchased from the ATCC organization while CAL-1 cells were a gift from Philippe Pierre laboratory (53). THP-1 NAP1-V84M^KI/WT^ cells and CAL-1 NAP1-V84M^KI/KI^ cells were generated for this study (see section below). All these cells were cultured at 37°C, 5% CO_2_, in Roswell Park Memorial Institute (RPMI) 1640 GlutaMAX^TM^ medium (Gibco-Life Technologies) supplemented with 10% FCS, 100 units/mL of Penicillin and 100 mg/mL of Streptomycin.

### Virus

The recombinant RABV-SAD (L16-B19) was previously described (28).

### Plasmids

All plasmids used in this study are listed in the following table. For each construction made in the pcDNA3.1 hygro(-) backbone the vector has been linearized using *NheI* and *NotI* restriction sites. Then, the inserts were introduced by Gibson technology using NEBuilder® HiFi DNA Assembly Master Mix (New England Biolabs, #E2621) according to the manufacturer’s instructions. The synthetic genes encoding TBK1, NAP1, SINTBAD, and TANK proteins were purchased from Twist Bioscience. All the oligonucleotides used for cloning were purchased from Eurofins Genomics.

Plasmids pEF6_ΔRIG-I(2xCARDs), pIFN-β-Luc and pcDNA3.1_IRF3-5D were previously described (30,54,55). The plasmid pRL-TK was purchased from Promega Corporation (#E2241). The plasmids psPAX2 (Addgene, 12260) and pMD2.G (Addgene, 12259) come from Didier Trono’s lab (EPFL, Lausanne, Switzerland). The lentiCRISPRv2 was a gift from Feng Zhang (Addgene plasmid # 52961; http://n2t.net/addgene:52961; RRID: Addgene_52961). The prK7_FLAG-STING antibody was a gift from Bo Zhong.

| Plasmid name | Gene | Tag | Mutation(s) | Origin |
| --- | --- | --- | --- | --- |
| **pcDNA3.1 hygro(-) backbone** |  |  |  |  |
| TBK1 | Human TBK1 | - | - | Ref (28) |
| TBK1 1-668 | Human TBK1 | - | ΔCTD (1-668) | This work |
| NAP1 | Human NAP1 | - | - | This work |
| NAP1 S34A | Human NAP1 | - | S34A | This work |
| NAP1 S82A-S83A | Human NAP1 | - | S82A-S83A | This work |
| NAP1 V84M | Human NAP1 | - | V84M | This work |
| NAP1 K109E | Human NAP1 | - | K109E | This work |
| NAP1 S120A | Human NAP1 | - | S120A | This work |
| NAP1 S206A | Human NAP1 | - | S206A | This work |
| NAP1 P302H | Human NAP1 | - | P302H | This work |
| NAP1 S318A | Human NAP1 | - | S318A | This work |
| NAP1 T320A | Human NAP1 | - | T320A | This work |
| NAP1 S343A | Human NAP1 | - | S343A | This work |
| NAP1 S353A | Human NAP1 | - | S353A | This work |
| NAP1 6XA | Human NAP1 | - | S34A-S82A-S83A-S120A-S318A-S343A | This work |
| NAP1 8XA | Human NAP1 | - | S34A-S82A-S83A-S120A-S206A-S318A-S343A-S353A | This work |
| SINTBAD | Human SINTBAD | - | - | This work |
| TANK | Human TANK | - | - | This work |
| IRF3-5D | Human IRF3 | - | S396D, S398D, S402D, T404D and S405D | Ref (30) |
| GFP | - | - | - | This work |
| GFP-TBK1 | Human TBK1 | GFP (N-ter) | - | Ref (28) |
| GFP-TBK1 1-668 | Human TBK1 | GFP (N-ter) | ΔCTD (1-668) | This work |
| GFP-TBK1 K38A | Human TBK1 | GFP (N-ter) | K38A | This work |
| GFP-TBK1 S172A | Human TBK1 | GFP (N-ter) | S172A | Ref (28) |
| GFP-TBK1 Y179A | Human TBK1 | GFP (N-ter) | Y179A | Ref (28) |
| GFP-NAP1 | Human NAP1 | GFP (N-ter) | - | This work |
| GFP-PP2A-Cα | Human PP2A (catalytic subunit) | GFP (N-ter) | | This work |
| GFP-SINTBAD | Human SINTBAD | GFP (N-ter) | - | This work |
| GFP-SINTBAD 1-280 | Human SINTBAD | GFP (N-ter) | ΔCTD (1-280) | This work |
| GFP-SINTBAD 280-615 | Human SINTBAD | GFP (N-ter) | ΔNTD (280-615) | This work |
| GFP-TANK | Human TANK | GFP (N-ter) | - | This work |
| GFP-TANK 1-129 | Human TANK | GFP (N-ter) | ΔCTD (1-129) | This work |
| GFP-TANK 129-425 | Human TANK | GFP (N-ter) | ΔNTD (129-425) | This work |
| mC | - | - | - | This work |
| mC-TBK1 | Human TBK1 | mC (N-ter) | - | This work |
| mC-NAP1 | Human NAP1 | mC (N-ter) | - | Ref (28) |
| mC-NAP1 1-211 | Human NAP1 | mC (N-ter) | ΔCTD (1-211) | This work |
| mC-NAP1 210-392 | Human NAP1 | mC (N-ter) | ΔNTD (210-392) | This work |
| mC-SINTBAD | Human SINTBAD | mC (N-ter) | - | This work |
| mC-SINTBAD 1-280 | Human SINTBAD | mC (N-ter) | ΔCTD (1-280) | This work |
| mC-SINTBAD 280-615 | Human SINTBAD | mC (N-ter) | ΔNTD (280-615) | This work |
| mC-TANK | Human TANK | mC (N-ter) | - | This work |
| mC-TANK 1-129 | Human TANK | mC (N-ter) | ΔCTD (1-129) | This work |
| mC-TANK 129-425 | Human TANK | mC (N-ter) | ΔNTD (129-425) | This work |
| MYC-NAP1 | Human NAP1 | MYC (N-ter) | - | This work |
| FLAG-TBK1 | Human TBK1 | FLAG (N-ter) | - | This work |
| FLAG TBK1 1-668 | Human TBK1 | FLAG (N-ter) | ΔCTD (1-668) | This work |
| FLAG-SINTBAD | Human SINTBAD | FLAG (N-ter) | - | This work |
| HA-SINTBAD | Human SINTBAD | HA (N-ter) | - | This work |
| HA-TANK | Human TANK | HA (N-ter) | - | This work |
| V5-TANK | Human TANK | V5 (N-ter) | - | This work |
| **Others backbones** |  |  |  |  |
| pEF6_ΔRIG-I(2xCARDS) | Human RIG-I | - | ΔNTD (285-925) | Ref (55) |
| pRL-TK | Renilla Luc | - | - | Promega, #E2241 |
| pIFN-β-Luc | Firefly Luc | - | - | Ref (54) |
| psPAX2 | HIV-1-Gag and Pol | - | - | Addgene, #12260 |
| pMD2.G | VSV-G | - | - | Addgene, #12259 |
| pLentiCRISPRv2_gRNA-TBK1 | Cas9 | - | Insertion of gRNA against *TBK1* | This work |
| pLentiCRISPRv2_gRNA-NAP1 | Cas9 | - | Insertion of gRNA against *NAP1* | This work |
| pLentiCRISPRv2_gRNA-SINTBAD | Cas9 | - | Insertion of gRNA against *SINTBAD* | This work |
| pLentiCRISPRv2_gRNA-TANK | Cas9 | - | Insertion of gRNA against *TANK* | This work |
| pRK7_FLAG-STING | Human STING | FLAG (N-ter) |  | Gift from Bo Zhong |

### Antibodies

Primary antibodies used in this study included mouse monoclonal **anti-Actin-β** (Millipore, #MAB1501) used at a 1:10 000 dilution for WB analysis; mouse monoclonal **anti-α-Tubulin** (Sigma, #T6199) used at a 1:1000 dilution for WB analysis; mouse monoclonal **anti-FLAG** (Cell Signaling Technology, #8146) used at a 1:1000 dilution for WB analysis, at a dilution 1:100 for IF analysis and at a 1:100 dilution for IP analysis ; rabbit monoclonal **anti-FLAG** (Cell Signaling Technology, #14793) used at a 1:1000 dilution for WB analysis and at a 1:100 dilution for IF analysis; rabbit monoclonal **anti-GAPDH** (Cell Signaling Technology, #2118) used at a 1:1000 dilution for WB analysis; mouse monoclonal **anti-GFP** (Roche, #11814460001) used at a 1:1000 dilution for WB analysis and a 1:50 dilution for IP analysis; monoclonal mouse **anti-HA** (Abcam, #ab130275) used at a 1:100 dilution for IP analysis ; rabbit polyclonal **anti-IRF3** (Santa Cruz, #sc-9082) used at a 1:1000 for WB analysis and a 1:100 dilution for IF analysis ; rabbit polyclonal **anti-MAVS** (Bethyl, #A300-782A) used at a 1:1000 dilution for WB analysis ; rabbit monoclonal **anti-mCherry** (Cell Signalin Technology, #43590) used at a 1:1000 dilution for WB analysis ; rabbit monoclonal **anti-MYC** (Cell Signaling Technology, #2278) used at a 1:1000 dilution for WB analysis ; mouse monoclonal **anti-RABV-N** (#81C4, previously described(56)) used at a 1:2500 dilution for IF analysis ; rabbit monoclonal **anti-NAP1** (Abcam, #192253) used at a 1:1000 dilution for WB analysis and a 1:100 dilution for IF analysis ; rabbit monoclonal **anti-NAP1** (Proteintech, #15042-1-AP) used at a 1:1000 dilution for WB analysis and a 1:900 dilution for IF analysis ; mouse monoclonal **anti-RABV-P** (#26G6, previously described (57) used at a 1:1000 dilution for WB analysis ; rat monoclonal **anti-PP2A-Aα** (Santa Cruz ; #sc-56953) used at a 1:100 dilution for IF analysis, mouse monoclonal **anti-RIG-I** (Santa Cruz, #sc-376845) used at a 1:1000 dilution for WB analysis ; rabbit monoclonal **anti-SINTBAD** (Cell Signaling Technology, #8605) used at a 1:1000 dilution for WB analysis ; rabbit polyclonal **anti-SINTBAD** (Abcam, #ab106394) used at a 1:200 dilution for IF analysis ; mouse monoclonal **anti-TANK** (Santa Cruz, #sc-166642) used at a 1:1000 dilution for WB analysis and a 1:100 dilution for IF analysis ; rabbit monoclonal **anti-TBK1** (Abcam, #ab40676) used at a 1:1000 dilution for WB analysis and a 1:250 dilution for IF analysis ; rabbit monoclonal **anti-phospho-TBK1 (Ser172)** (Cell Signaling Technology, #5483) used at a 1:1000 dilution for WB analysis and a 1:250 dilution for IF analysis and mouse monoclonal **anti-V5** (Invitrogen, #R960-25) used at a 1:5000 dilution for WB analysis.

Secondary antibodies used for IF analysis in this study were purchased from Invitrogen and included Alexa 488 goat anti-rabbit IgG (#A-11008); Alexa 488 goat anti-mouse IgG (#A-11029); Alexa 488 goat anti-rat IgG (#A11006); Alexa 568 goat anti-mouse IgG (#A-11031) and Alexa 647 goat anti-rabbit IgG (#A-21246).

Secondary antibodies used for WB analysis in this study included anti-rabbit 800-conjugated IgG (Cell Signaling Technology, #5151), anti-mouse 680-conjugated IgG (Cell Signaling Technology, #5470), goat anti-rabbit IgG-HRP (Invitrogen, #31460) and goat anti-mouse IgG-HRP (Sigma, #A4416).

### Cell treatment with reagents

To specifically trigger the production of type I IFN through the cGAS-STING pathway, cells were treated with diABZI (Invitrogen, #tlrl-diabzi-2), a STING agonist. On the stimulation day, the cell medium is replaced with a medium containing diABZI at a concentration of 100 ng/mL for 6 to 24 h. To assess the role of PP2A phosphatases, cells were treated with LB-100 (Abcam, # ab285402), a potent and specific inhibitor of the PP2A family. On the inhibition day, the cell medium is replaced by a medium containing LB-100 at a concentration of 5 µM for 4h. To differentiate THP-1 cells into macrophage-like cells, they were treated with Phorbol 12-myristate 13-acetate (PMA, ThermoFisher Scientific, #15476069) at a concentration of 100 ng/mL for 48h.

### Cell transfection

Transfections with plasmids (see list above) were conducted using either Lipofectamine 2000 for HEK293T cells (ThermoFisher, #11668019) or Lipofectamine 3000 for HAP1 cells (ThermoFisher, # L3000015), according to the manufacturer’s instructions. One day prior to transfection, cells were seeded in a 12 or 24-well plate and incubated at 37°C - 5% CO_2_. After 24 h of growth, the culture medium was replaced by DMEM or IMDM supplemented with 2.5% FCS (without antibiotics). Next, the plasmids were dispersed in opti-MEM (Gibco, #31985–047) and the transfection reagent was added. After incubating for 5 min at room temperature (RT), the transfection mixes were added to the cells. Subsequently, the cells were incubated at 37°C - 5% CO_2_ and analyzed up to 36h p.t. In 12-well (resp. 24-well) plates up to 1µg (resp. 0.5µg) of plasmids is transfected.

The 5’ppp-dsRNA (InvivoGen, #tlrl-3prna) was used at the final 4 mg/mL concentration. HEK293T cells were transfected using Lyovec (InvivoGen, #lyec-1) while HAP1 cells were transfected with Lipofectamine 3000 according to the manufacturer’s instructions and with the procedure described above. The cells were then analyzed 6h p.t.

### Generation of CRISPR-Cas9 KO HEK293T cells

HEK293T KO cells were generated via lentiviral transduction using single guide RNA (sgRNA) oligonucleotide sequences cloned into the lentiCRISPR v2 plasmid system (58). Briefly, the plasmid lentiCRISPRv2 was linearized using the *BsmB*I restriction site and sgRNA duplexes were inserted inside it. The following sequences were used to target the specific gene:

*TBK1*: 5’-CACCGAAGAACCTTCTAATGCCTA-3’ (exon 4),
*AZI2*: 5’-CACCGGCCTATCATGCATATCGAG-3’ (exon 3),
*SINTBAD*: 5’-CACCGCATCATCCGGGTACCCGAC-3’ (exon 9)
*TANK*: 5’-CACCGAGCCTTCCGGCAGGCATGCA-3’ (exon 2).

A non-targeting sequence 5’-GTTCCGCGTTACATTAACTTA-3’ was used as negative control (referred to as CTRL) and has been previously described (59).

To obtain stocks of lentiviral particles, HEK293T cells were with 0.9 µg of lentiCRISPR v2 plasmid expressing sgRNA targeting an exon within the gene of interest (*i.e. TBK1*, *AZI2*, *SINTBAD* or *TANK*), using Lipofectamine 2000, along with 1.4 µg of psPAX2 and 0.45 µg of pMD2.G. One day p.t., the culture medium was removed, and cells were cultured for an additional 24 hours in fresh medium. Supernatants of the culture were then collected, filtered (0.45 µm), and directly used to transduce new HEK293T cells by spinoculation (centrifugation at 800g for 30 min. at 32°C). Three days after transduction, cells were selected by the addition of 0.75 µg/mL of puromycin in the culture medium. At the end of the selection, a limiting dilution procedure was performed to isolate single clones. Then, the clones were screened and the KO of target genes was checked by WB.

### Generation of KI THP-1 and CAL-1 cells

To introduce the *AZI2* p.V84M mutation in THP1 and CAL1 cell lines, the CRISPR-Cas9 system combined with homology-directed repair (HDR) was employed. sgRNAs targeting the *AZI2* locus were designed using the IDT CRISPR-Cas9 guide RNA design tool. The selected sgRNA was synthesized as an Alt-R CRISPR-Cas9 crRNA and combined with the Alt-R tracrRNA to form a functional ribonucleoprotein (RNP) complex. The RNP complex was assembled with Alt-R S.p. HiFi Cas9 Nuclease V3 (IDT) to enhance specificity and reduce off-target effects. A single-stranded DNA (ssDNA) HDR template containing the desired p.V84M mutation, along with homologous flanking sequences of approximately 100 bp on each side, was designed and synthesized (IDT). The ssDNA donor was co-delivered with the RNP complex into THP1 and CAL1 cells via electroporation using the Lonza Nucleofector 2AD system with cell line-specific transfection conditions. After electroporation, cells were allowed to recover for 48 hours before single-cell sorting was performed using a Sony MA900 cell sorter. Single cells were plated into 96-well plates containing RPMI-1640 medium supplemented with 10% fetal bovine serum (FBS) and 1% penicillin-streptomycin. Clones were expanded for genomic DNA extraction. Genomic DNA from expanded clones was extracted. The AZI2 target region was amplified by PCR, and Sanger sequencing was performed to identify wild-type (WT) and knock-in (KI) clones (heterozygous or homozygous). Sequence alignment and mutation verification were carried out using APE software. Successfully edited clones carrying the p.V84M mutation were further expanded and characterized for downstream functional assays as well as some WT clones to perform experiments with cells that have undergone the whole process but remained WT.

### Viral infection

Cell monolayers (confluency between 60 to 80%) were infected with recombinant RABV-SAD (L16-B19) diluted in DMEM (or IMDM) at a multiplicity of infection (MOI) of 3. After 1h of adsorption at RT, the inoculum was removed and the cells were maintained in DMEM (or IMDM) supplemented with 2.5% FCS and then incubated at 37°C - 5% CO_2_. For « mock » conditions, cells were incubated for 1h at RT only with DMEM (or IMDM).

### Luciferase reporter assay

HEK293T cells were seeded in 24-well plates at a cell density of 1.10^5^ per well and transfected (total of 650 ng of plasmids per well) the next day using lipofectamine 2000. In all experiments, cells were transfected with 25 ng of pRL-TK plasmid encoding the Renilla Luc (used as internal control) and with 125 ng of pIFN-β-Luc plasmid encoding Firefly Luc under the control of IFN-β promoter. Additionally, 500ng of specific plasmids required for the dedicated experiment were also transfected. At the indicated time p.t., cell extracts were prepared using Passive cell Lysis Buffer (Promega, #E1910) according to the manufacturer’s instructions and were then subjected to WB analysis or Luc reporter gene activity using the Dual-Luciferase® Reporter Assay System (Promega, #E1910). Each experiment was performed in triplicate. For each sample, the Firefly Luc activity were normalized to Renilla Luc activity, and the mean fold induction of Firefly Luc was compared to that in non-stimulated conditions.

### RT-qPCR to measure *IFN-β* mRNA

Control and KO HEK293T (or HAP1) cells were infected with RABV for 16 h or transfected for 24 h to induce the IFN induction pathway. Cells were lysed and total RNAs were isolated and purified with the NucleoSpin RNA kit (Macherey-Nagel, #740955) following the manufacturer’s instructions. The purity and concentrations of extracted RNAs were assessed spectroscopically using a Nanodrop system. Then 500 ng of total RNAs were used to synthesize complementary DNAs (cDNAs) using the High Capacity cDNA Reverse Transcription Kit (ThermoFischer, #4368814). Human IFN-β and GAPDH cDNAs were quantified by qPCR using the Power SYBR Green Master Mix (ThermoFischer, #4309155) and specific primers (sequences below). The 2^−ΔCt^ method was used to quantify the relative amount of *IFN-β* mRNA to *GAPDH* mRNA for each condition and the results were presented as fold induction relative to the control condition. The following primers have been used to amplify the IFN-β cDNA (For: 5’- GTCTCCTCCAAATTGCTCTC-3’ / Rev: 5’-ACAGGAGCTTCTCACACTGA-3’) and the GAPDH cDNA (For: 5’-AACAGCCTCAAGATCATCAGCAA-3’ / Rev: 5’-TCTTCTGGGTGGCAGTGAT-3’).

### WB analysis

The cell monolayers were washed with cold PBS and lysed on ice for 30 min using a RIPA lysis buffer (50 mM Tris-HCl pH 7.4, 100 mM NaCl, 0.5% NP-40) supplemented with 1 mM PMSF, a protease inhibitors cocktail (Roche, #11-836-170-001) and NaF (Sigma, #67414). Lysates were then clarified at 16,000g for 10 min to remove the insoluble fraction. Eventually, protein concentrations of samples were measured using Bradford assays (Bio-Rad, #5000006), following the manufacturer’s instructions. Lysates were then diluted in 1× reducing Laemmli buffer (240 mM Tris-HCl pH6.8, 8% SDS, 400 mM DTT, 40% Glycerol 0.4% bromophenol blue). Samples were boiled at 95°C for 10 min and separated by SDS-PAGE. Proteins were transferred onto nitrocellulose membranes (0.2 µm) and saturated in TBS - 0.1% Tween 20 containing 5% BSA for 1 h at RT. Membranes were then incubated overnight at 4°C with the specific primary antibody in TBS supplemented with 0.1% Tween 20 and 1% BSA. After extensive washing, membranes were incubated for 1 h at RT with either Fluor-conjugated IgG secondary antibodies or horseradish peroxidase-conjugated secondary antibodies. After washing, the membranes were scanned with the Odyssey infrared imaging system (LI-COR, Lincoln, NE).

### Immunofluorescence analysis

HEK293T, HAP1 and THP-1 cells were directly seeded on glass coverslips in 12-well plates, while for CAL-1 cells the glass coverslips were first coated with poly-D-Lysine (50% [vol/vol] in sterile water) for 30 min before cell seeding. To allow THP-1 cells to adhere to glass coverslips, just after the seeding, they were differentiated into macrophage-like cells.

Cell monolayers were washed with PBS and then fixed using one of the following methods to optimize the visualization of the condensates (28). Cells were either fixed for 15 min at RT with 4% paraformaldehyde (PFA) and further permeabilized for 5 min with 0.1% Triton X-100 in PBS, or fixed for 10 min at -20°C with a methanol/acetone solution (37.5%/50% [v/v] respectively) before rehydration for 3 min in PBS. Following saturation with 1% BSA in PBS for 15 min, cells were incubated with the indicated primary antibodies for 1h at RT. Then, cells were washed three times with PBS, and incubated for 45 min at RT with the appropriate Alexa fluor conjugated secondary antibodies. After washing, the nuclei were counterstained with 0.1 mg/mL of DAPI in PBS for 5 min. Coverslips were mounted with Immuno-mount (ThermoFisher). Images were sequentially acquired using a laser scanning SP8 confocal microscope (Leica) with a 63x oil immersion objective (HC Plan Apo, NA: 1.4, Leica). We used 405, 488, 561, and 647 nm wavelengths to excite DAPI, GFP/AF488, mCherry/AF561, and AF647 respectively and their fluorescence was collected through the following bandpass 415-455, 498-538, 571-611 and 657-697 nm. All images shown in the study are representative of at least three independent experiments. Images were resized, organized, and labeled using Fiji-ImageJ software or LAS AF Lite.

### Co-immunoprecipitation assays

HEK293T cells were seeded in 12-well plates. 24 h p.t. with plasmids encoding the indicated tagged proteins, cells were lysed as indicated in the section “WB analysis”. Immunoprecipitations were performed by incubating whole cell lysates with mouse monoclonal anti-FLAG, anti-GFP, or anti-HA antibodies for 2h at 4°C under gentle rotation. Pierce^TM^ protein G-magnetic beads (ThermoFisher, #88848) were added for 2h at 4°C under gentle rotation. The beads were washed 3 times with RIPA buffer, and protein complexes were eluted in 2x reducing Laemmli buffer at 95°C for 10 min, then subjected to SDS-PAGE and WB.

### Tryptic digestion of NAP1 protein bands for mass spectrometry analysis

HEK293T cells (1.2 × 10^6^ cells) were transfected with plasmids encoding GFP-NAP1 or co-transfected with plasmids encoding GFP-NAP1 and mC-TBK1. After 24 hours, NAP1 was immunoprecipitated using an anti-GFP antibody and the immunoprecipitated proteins were separated by SDS-PAGE. Gel bands were stained with Instant-Blue^TM^ Coomassie and subsequently excised for mass spectrometry analysis. Briefly, the bands of interest were destained three times by incubation for 20 min under agitation in a solution of 50 mM ammonium bicarbonate and 50% acetonitrile. They were then completely dehydrated in 100% acetonitrile for 10 min, followed by a vacuum centrifugation step. Trypsin digestion was performed overnight at 37°C under agitation using trypsin gold mass Spectrometry grade (Promega, #V5280). Digestion was stopped during the two extraction steps by incubating the gel bands in 1% formic acid for 30 min. Finally, extracted samples were evaporated using a vacuum centrifuge and stored at -20°C until analysis.

### Liquid chromatography-mass spectrometry analysis of tryptic peptides

Peptides generated by in-gel trypsin digestion were dried, resuspended in loading buffer (2% acetonitrile and 0.05% trifluoroacetic acid in water), and analyzed by nanoLC-MS/MS using a nanoElute liquid chromatography system (Bruker) coupled to a timsTOF Pro 2 mass spectrometer (Bruker). Briefly, peptides were desalted on a C18 reverse-phase TRAP cartridge (C18 Acclaim Pepmap100, 5 µm, 5 mm length) using loading buffer and then eluted at a flow rate of 400 nL/min from an Aurora 3 analytical column (ION OPTIK, 25 cm × 75 µm ID, 1.7 µm, C18) with a gradient of 0–35% solvent B over 30 min. Solvent A consisted of 0.1% formic acid and 2% acetonitrile in water, while solvent B was 99.9% acetonitrile with 0.1% formic acid. MS and MS/MS spectra were recorded from m/z 100 to 1700, with an ion mobility scan range of 0.6 to 1.5 V.s/cm². MS/MS spectra were acquired in PASEF (Parallel Accumulation – Serial Fragmentation) mode, with 10 PASEF MSMS scans.

### Mass spectrometry data analysis

A database search was performed using the Mascot search engine (version 2.6., Matrix Science, London, UK) against a database containing amino acid sequences of GFP-NAP1 and mC-TBK1 proteins. Database searches were conducted with semi-trypsin cleavage specificity and up to four miscleavages. Oxidation of methionine and phosphorylation of serine, threonine, and tyrosine were set as variable modifications. Peptide and fragment tolerances were set at 15 ppm and 0.05 Da, respectively. The mass spectrometry proteomics data have been deposited to the ProteomeXchange Consortium via the PRIDE partner repository with the dataset identifier PXD062298 and 10.6019/PXD062298

### Live imaging microscopy

For live imaging, cells were seeded in a 35 mm Glass Bottom µ-Dish (IBIDI, #81218) or in a 2.5cm² glass-bottom µ-slide (IBIDI, #80427). After 24h of growth, cells were transfected with plasmids encoding fluorescent proteins, as indicated in figure legends. From 18h p.t., the cell medium was replaced by FluoroBrite™ DMEM (Gibco-Life Technologies, #A1896702) supplemented with 10% FCS, then cells were incubated at 37°C - 5% CO_2_ and data acquisition started on a confocal spinning-disk microscope.

Image acquisition was performed on an inverted Nikon Ti Eclipse microscope coupled with a spinning disk (Yokogawa CSU-X1) and a 100X objective (Apochromat TIRF oil immersion, NA: 1.49). A blue laser (488 nm, Vortan, 150 mW) was used for the excitation of GFP-fused proteins and a yellow laser (561nm, Coherent, 100 mW) for the excitation of mCherry-fused proteins. A quad-band dichroic mirror (405/491/561/641 nm, Chroma) and band-pass filters of 525/45 nm (Semrock), 607/36 nm (Semrock) and 692/40 nm (Semrock) were used respectively to ensure specific detection of the fluorescence. Images were recorded with a Prime 95B sCMOS camera (Photometrics). FRAP experiments were performed using iLas 2module (GATACA systems) and the whole system was driven by MetaMorph software version 7.7 (Molecular Devices).

### Characterization of liquid condensates in live imaging

To assess the liquid properties of the condensates, cells were treated with 1,6-HD (Sigma, #240117) or subjected to a hypotonic shock. For 1,6-HD treatment, during the acquisition, culture medium containing 1,6-HD at a 2X concentration was directly added to the cells to reach the working concentration of 10%. Data acquisition was maintained until the complete disappearance of condensates (∼15 min). For the hypotonic shock, during the acquisition, 3 volumes of sterile water were directly added to the cells. Data acquisition was maintained until the condensates reappeared (∼15 min) following their complete disappearance (∼5 min).

### FRAP experiments

FRAP experiments were performed on HEK293T cells expressing GFP-NAP1, GFP-SINTBAD, GFP-TANK or GFP-TBK1 (along with mC-NAP1 for this latter) for 20 h, and acquired using the confocal spinning-disk microscope described above.

For SINTBAD and TANK condensates, FRAP experiments were performed using the following sequences: 5 s of pre-bleach (frame rate of 2 images per second), 40ms to bleach GFP-SINTBAD and GFP-TANK, 10 s of a first post-bleach sequence (frame rate of 5 images per second) and 20 s of a second post-bleach sequence (frame rate of 2 images per second). Alternatively, FRAP experiments on GFP-NAP1 and GFP-TBK1 (inside mC-NAP1 condensates) were performed using the following sequence: 5 s of pre-bleach (frame rate of 2 images per second), 40ms to bleach GFP-NAP1 and GFP-TBK1 chimeras, 10 s of a first post-bleach sequence (frame rate of 5 images per second) and 40 s of a second post-bleach sequence (frame rate of 2 images per second). In all cases, bleaching was performed in circular regions to bleach the whole condensate.

For the FRAP data analysis, we used homemade macros on Fiji-ImageJ software and Python performing the following steps. On every slide, the background was estimated and removed by measuring a region in the cytoplasm of the cell. To correct acquisitional bleaching, the mean fluorescence within the bleached region was normalized by the mean value of a neighboring condensate (in the same cell when possible). Then a second normalization was applied by dividing the results by the mean value of the pre-bleach sequence to allow comparison of recovery curves.

### Quantifications

The proportion of cells transfected with 5’ppp-dsRNA presenting condensates (TBK1, NAP1, SINTBAD or TANK) was determined by counting at least 30 randomly selected cells in each condition from three independent experiments. It was assumed that analyzed micrographs corresponded to median sections of cells.

Similarly, the proportion of transfected cells presenting an IRF3 nuclear localization was determined by counting at least 30 randomly selected cells per condition from three independent experiments.

To determine the critical concentration at which different mutants of NAP1 lead to LLPS, we measured their areal fluorescence in the dilute phase (*i.e.* in the cytosol) of transfected cells presenting condensates from confocal images. For this, we applied the method outlined in (60) and previously employed by our group (61), by using Fiji/ImageJ software. We assumed that the local concentration of NAP1 is proportional to its local fluorescence intensity per pixel. Initially, we established the background intensity by measuring the areal fluorescence intensity in a 50×50 pixel area outside of cells. We then randomly selected five representative areas (each 21×21 pixels) of the dilute phase and calculated their mean fluorescence intensity. To determine the concentration of NAP1 in the dilute phase, the background value was subtracted from the areal fluorescence in the dilute phase. For each condition, 20 randomly selected cells were quantified from three independent experiments.

### Statistical analysis

All presented data come from at least three independent experiments. Data are expressed as mean ± standard deviation (SD) and statistical analysis was performed using Prism (GraphPad 10 software). All experimental values are shown on the corresponding graph. Data comparisons were performed using the Student t-test, ANOVA test or Kolmogorov-Smirnov test. Statistical details of each experiment are indicated in the figure legends.

## Acknowledgements

This work was supported by the « Prix Bettencourt Coups d’élan pour la recherche française » attributed to YG, a grant from the Agence Nationale de la Recherche, France, (ANR-24-CE15-7522, ChaConAdapt) to Y.G., and a grant from the Fondation pour la Recherche Médicale, France, to Y.G. (including a post-doctoral grant to D.G.) This work has benefited from the core facilities of Imagerie-Gif, supported by “France-BioImaging” (ANR-10-INBS-04-01) and from the facilities and expertise of the I2BC proteomics platform (Proteomique-Gif, SICaPS, supported by IBiSA, Ile de France Region, Plan Cancer, CNRS and Paris-Saclay University). We thank Bruker for technical assistance.

## Author contributions

Conceptualization, D.G., Q.R., F.R.-L., and Y.G.; methodology, D.G., Q.R., K.B., B.B.-M., M.-L.F., C.L.-G., F.R.-L, and Y.G.; investigations, D.G., Q.R., F.R., B.L., A.G., L.S., O.P., D.H.-N., A.L., and M.T.; formal analysis, D.G., Q.R., F.R., L.S., C.L.-G., F.R.-L., and Y.G.; supervision, B.B.-M., M.-L.F., Y.C., A.B., C.L.-G., F.R.-L., and Y.G.; funding acquisition, Y.G.; writing—original draft, D.G., and Y.G.; writing—review and editing, D.G., Q.R., C.L.-G., F.R.-L., and Y.G.

## Declaration of interests

The authors declare no competing interests.

## Supplemental figures

**Fig. S1.**
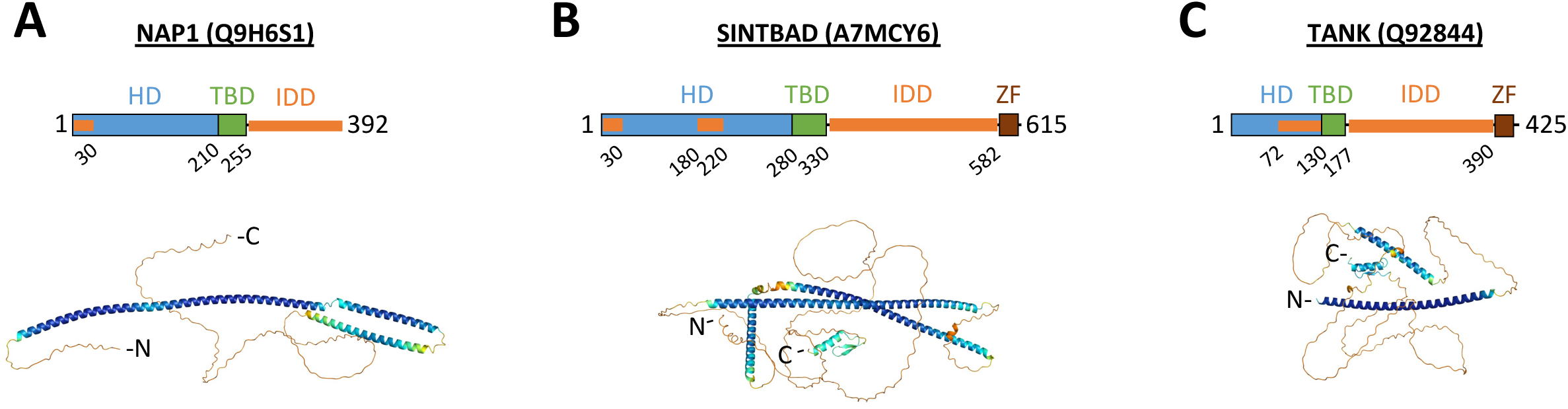
Modular organization of NAP1, SINTBAD and TANK. **A-C)** Bar diagram showing the domain organization of NAP1 (**A**), SINTBAD (**B**) and TANK (**C**). HD, homodimerization domain; IDD, intrinsically disordered domain; TBD, TBK1 binding domain; ZF, zinc finger. The Alphafold3 prediction of the structure of each protein is also presented. Putative intrinsically disordered regions were predicted based on Alphafold3 structures using the MobiDB software (62).

**Fig. S2.**
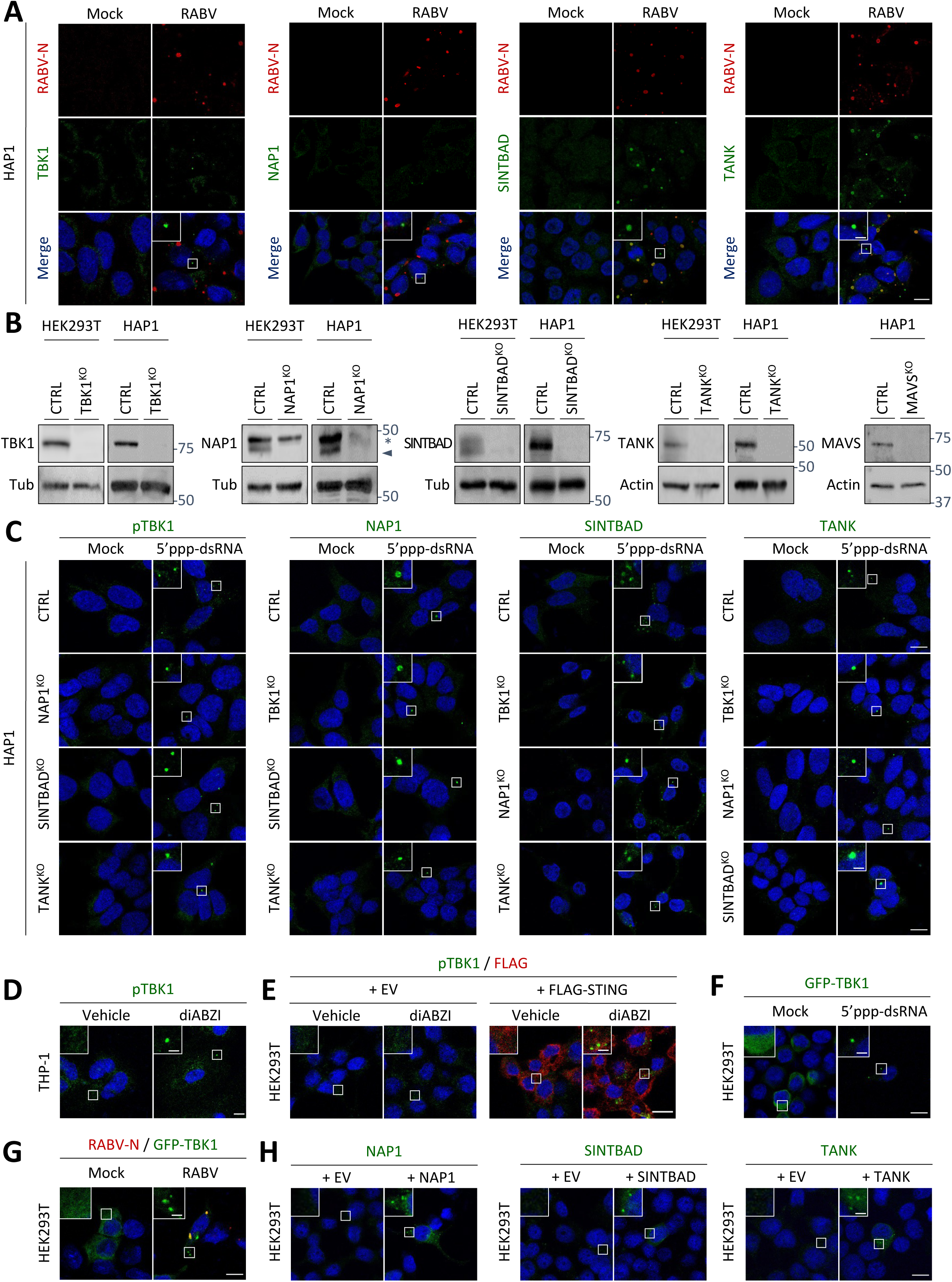
Detection of danger signals as well as ectopic expression induce condensation of TBK1, NAP1, SINTBAD and TANK. **A)** HAP1 cells were infected with RABV (MOI=3) or not (Mock). At 16 h p.i., cells were fixed before co-staining with anti-N and either anti-pTBK1, anti-NAP1, anti-SINTBAD or anti-TANK antibodies and analyzed by confocal microscopy. **B)** Parental (CTRL) and indicated KO HEK293T or HAP1 cells were lysed. Extracts were analyzed by WB with either anti-TBK1, anti-NAP1, anti-SINTBAD, anti-TANK or anti-MAVS antibody. Anti-Tubulin or anti-Actin antibodies were used as loading control. For the anti-NAP1 antibody, the star indicates an unspecific band whereas the arrow indicates the band corresponding to NAP1. **C)** Parental (CTRL) and KO HAP1 cells (TBK1^KO^, NAP1^KO^, SINTBAD^KO^ and TANK^KO^) were incubated with medium containing Lipofectamine 3000 alone (Mock) or transfected with 5’ppp-dsRNA using Lipofectamine 3000 (5’ppp-dsRNA). After 6 h, cells were fixed before staining with anti-pTBK1, anti-NAP1, anti-SINTBAD or anti-TANK antibody and analyzed by confocal microscopy. **D)** THP-1 cells (differentiated with PMA for 48 h) were treated with diABZI for 6 h. Cells were fixed before staining with anti-pTBK1 and anti-FLAG antibodies and analyzed by confocal microscopy. **E)** HEK293T cells were transfected with a plasmid encoding FLAG-STING or an empty vector (EV). After 24 h, cells were treated with diabZI for 6h. Cells were fixed before staining with anti-pTBK1 (S172) antibody and analyzed by confocal microscopy. **F)** HEK293T cells were transfected with plasmids encoding GFP-TBK1. After 18 h, cells were incubated with a medium containing Lipofectamine 3000 alone (Mock) or transfected with 5’ppp-dsRNA using Lipofectamine 3000 (5’ppp-dsRNA). After an additional 6h, cells were fixed and analyzed by confocal microscopy. **G)** HEK293T cells were infected (MOI=3) with RABV or not (Mock) and transfected with a plasmid encoding GFP-TBK1. At 16 h p.i., cells were fixed before staining with anti-N antibody and analyzed by confocal microscopy. **H)** HEK293T cells were transfected with plasmids encoding untagged versions of NAP1, SINTBAD or TANK (or EV). After 24 h, cells were fixed before staining with anti-NAP1, anti-SINTBAD or anti-TANK antibody and analyzed by confocal microscopy. On microscopy images, square boxes have been magnified (3X) at the top left. Scale bars: 10µm and 2 µm for insets.

**Fig. S3.**
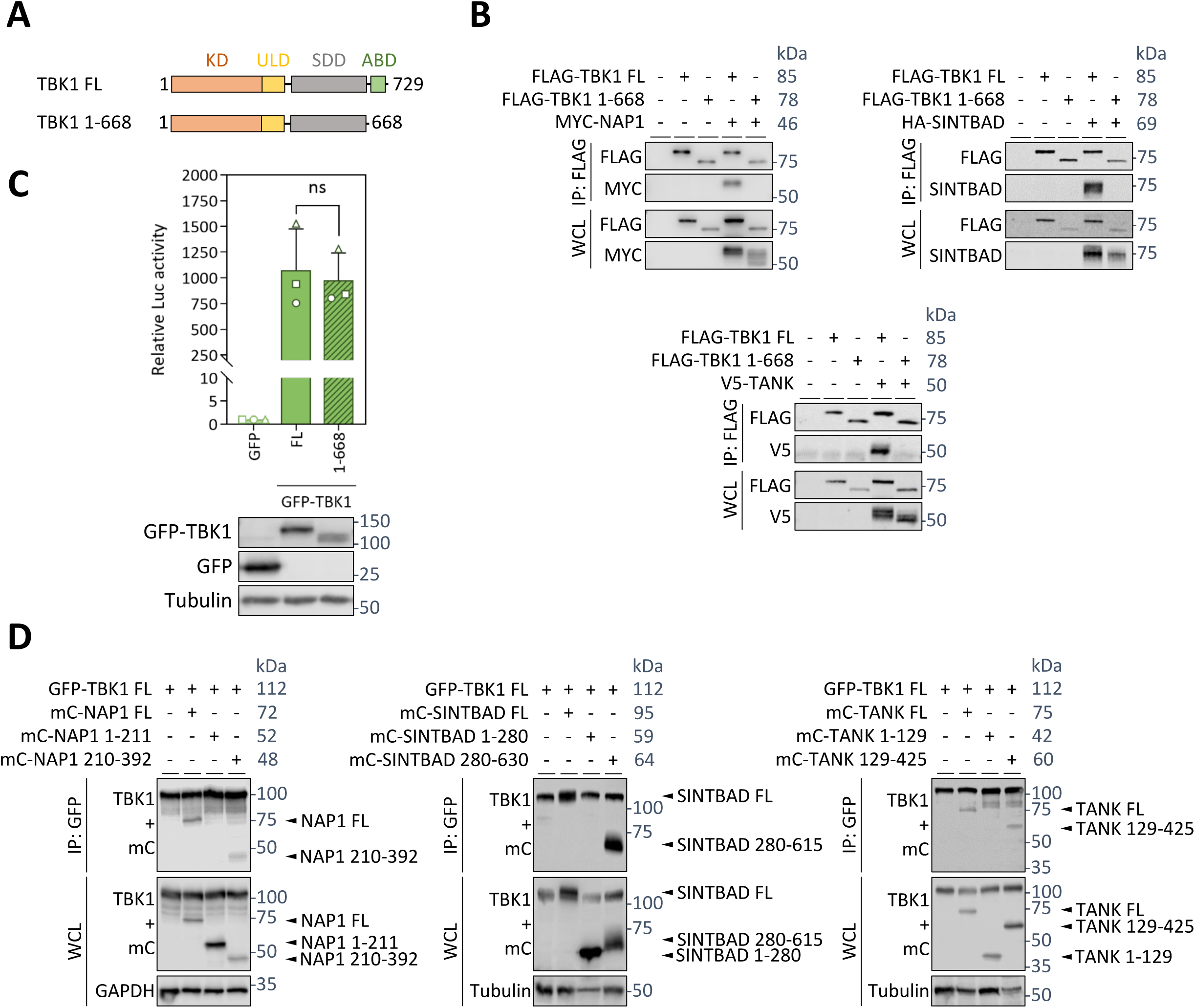
Characterization of TBK1 and TBK1-adaptor protein truncations. **A)** Bar diagram showing the domain organization of TBK1. The ABD domain is absent in deletion mutant TBK1 1-668. **B)** HEK293T cells were co-transfected with plasmids encoding FLAG-TBK1 (FL or 1-668) and MYC-NAP1 (upper left panel), HA-SINTBAD (upper right panel) or V5-TANK (bottom left panel) for 24 h. After immunoprecipitation of TBK1 (FL or 1-668) with an anti-FLAG antibody, cell lysates (WCL) and immunoprecipitated proteins (IP) were analyzed by WB with anti-FLAG and anti-MYC, anti-SINTBAD or anti-V5 antibodies. **C)** HEK293T cells were transfected with plasmids encoding GFP-TBK1 (FL or 1-668) or GFP alone and the plasmids of the Luc reporter assay. After 24 h, the induction of the IFN pathway was assessed using the Luc reporter assay (activation of the IFN-β promoter). Cell lysates were analyzed by WB using anti-GFP and anti-Tubulin antibodies. The bars represent the mean ± SD normalized to the GFP condition of 3 independent experiments. ns =non-significant (Student t-test). **D)** HEK293T cells were co-transfected with plasmids encoding GFP-TBK1 and indicated truncations of mC-NAP1 (left panel), mC-SINTBAD (middle panel) or mC-TANK (right panel). After immunoprecipitation of TBK1 with an anti-GFP antibody, cell lysates (WCL) and immunoprecipitated proteins (IP) were analyzed by WB with anti-TBK1, anti-mC and anti-GAPDH or anti-Tubulin antibodies. It is worth noting that mC-SINTBAD FL and GFP-TBK1 FL co-migrate in SDS-PAGE despite different molecular weights.

**Fig. S4.**
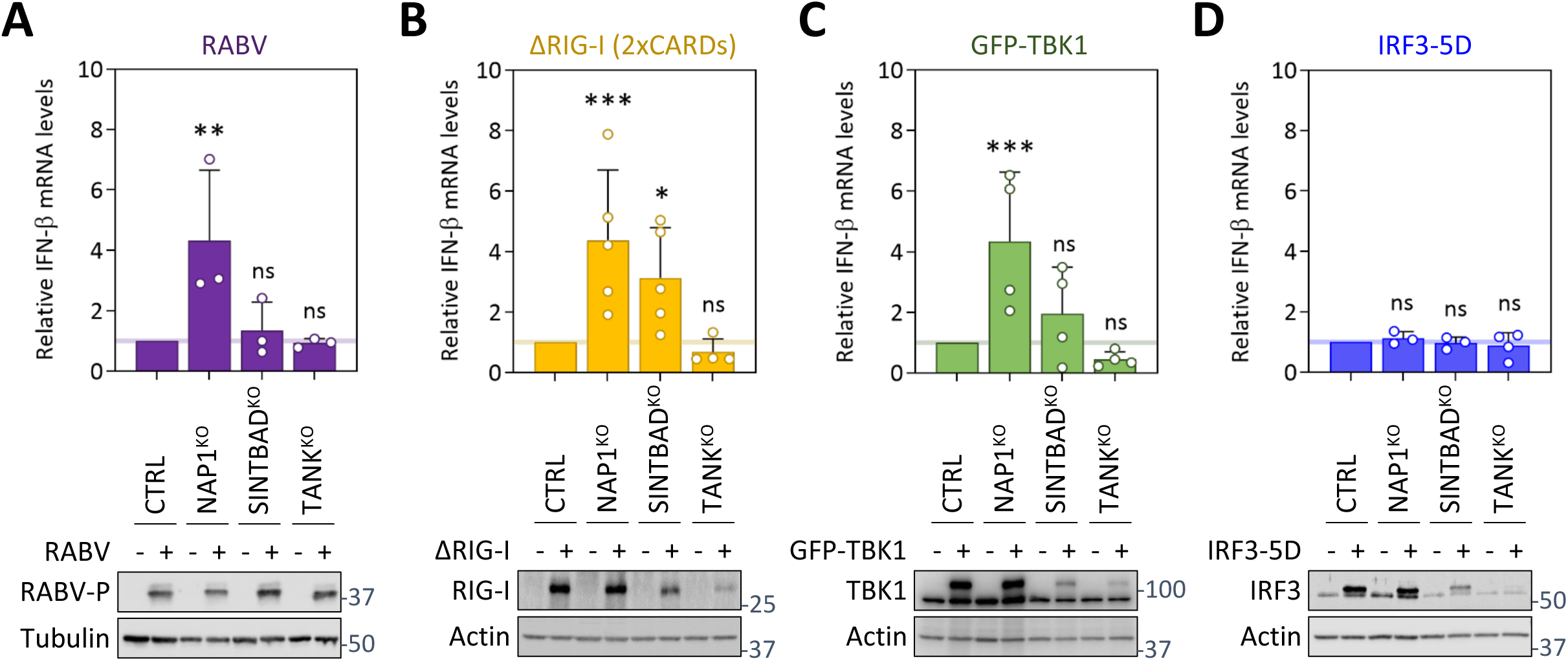
Impact of TBK1-adaptor proteins deficiency on IFN production in HAP1 cells. **A-D)** Parental (CTRL) and KO HAP1 cells (NAP1^KO^, SINTBAD^KO^, and TANK^KO^) were infected (MOI=3) with RABV (**A**), or transfected either with ΔRIG-I (**B**), GFP-TBK1 (**C**) or IRF3-5D (**D**) plasmids. At 16 h p.i. or 24 h p.t., cells were lysed and the relative levels of *IFN-β* mRNAs to *GAPDH* mRNAs were quantified by RT-qPCR. For each KO cell line, the *IFN-β* mRNA fold induction relative to the control cells is indicated. The bars represent the mean ± SD of at least three independent experiments. ns =non-significant; *P < 0.05; **P < 0.01, ***P < 0.001 (ANOVA test). Cell lysates were also analyzed by WB using anti-P, anti-RIG-I, anti-GFP, anti-IRF3, anti-Actin, or anti-Tubulin antibodies.

**Fig. S5.**
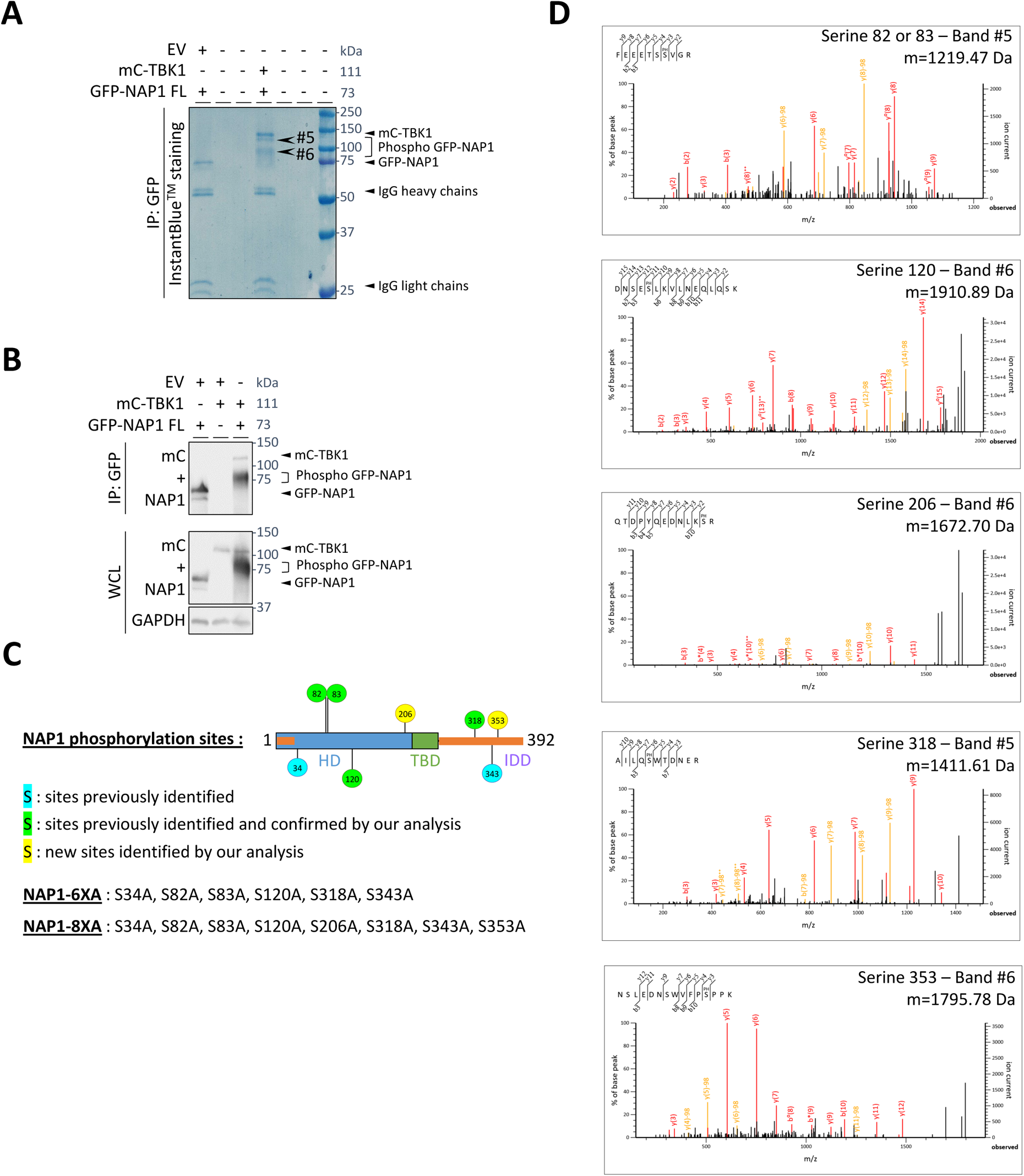
Identification of NAP1 phosphorylation sites by LC-MS. **A-B)** HEK293T cells were co-transfected with plasmids encoding GFP-NAP1 and mC-TBK1 (or EV) for 24 h. After cell lysis, GFP-NAP1 was immunoprecipitated using an anti-GFP antibody. **(A)** Immunoprecipitated proteins were separated by SDS-PAGE and stained by InstantBlue^TM^ Coomassie. **(B)** Cell lysates (WCL) and immunoprecipitated proteins (IP) were also analyzed by WB with anti-NAP1, anti-mC and anti-GAPDH antibodies. **C)** Scheme of NAP1 organization showing the position of phosphorylated serines identified in our analysis as either new phosphorylated sites (in yellow) or peptides previously identified in the literature (in green), or not identified in our analysis but previously reported (in blue).. Each site has been mutated in alanine and two multiple mutants have been constructed (NAP1 6XA and NAP1 8XA) as indicated. **D)** MS/MS spectra of phosphorylated peptides detected in digests of bands #5 and #6 of phosphorylated GFP-NAP1 labeled on SDS-PAGE (**A**) identify phosphorylation sites S82 or S83, S120, S206, S318, and S353 on NAP1. The mass of the phosphorylated peptide is indicated. Ion peaks labeled with ^++^ have two positive charges, those labeled with ° are dehydrated peptides, those labeled with * are deaminated peptides.

**Fig. S6.**
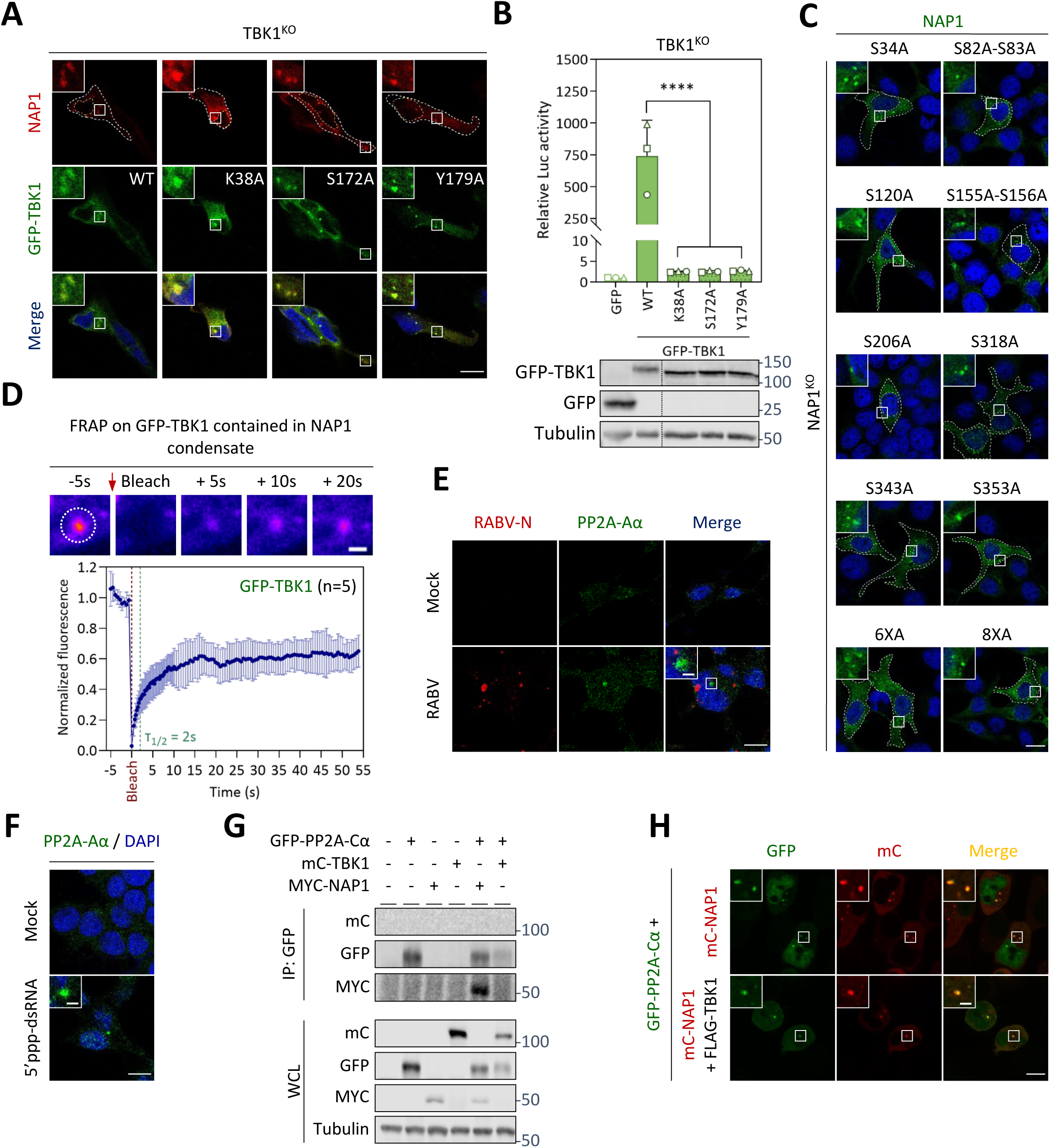
Characterization of TBK1 and NAP1 mutants as well as the accumulation of PP2A phosphatase complex inside NAP1 condensates. **A)** TBK1^KO^ HEK293T cells were co-transfected with plasmids encoding NAP1 and GFP-TBK1 WT or mutants (K38A, S172A or Y179A). After 24 h, cells were fixed before staining with an anti-NAP1 antibody and analyzed by confocal microscopy. **B)** TBK1^KO^ HEK293T cells were transfected with plasmids encoding GFP-TBK1 WT or mutants (K38A, S172A or Y179A) or GFP alone, and the plasmids of the Luc reporter assay. After 24 h, the induction of the IFN pathway was assessed using the Luc reporter assay (activation of the IFN-β promoter). Cell lysates were analyzed by WB using anti-GFP and anti-Tubulin antibodies. The bars represent the mean ± SD of three independent experiments and the corresponding data points are shown. ****P < 0.0001 (Anova). **C)** NAP1^KO^ HEK293T cells were transfected with plasmids encoding indicated NAP1 phosphorylation mutants. After 24 h, cells were fixed before staining with an anti-NAP1 antibody. **D)** HEK293T cells were co-transfected with plasmids encoding GFP-TBK1 and mC-NAP1. After 20 h, whole condensates were photobleached and the recovery of fluorescence was followed. by a confocal spinning-disk microscope. A representative FRAP experiment is shown and the photobleached region (Ø=3 µm) is indicated. The mean curve and the SD of 5 FRAP experiments is shown. **E)** HEK293T cells were infected (MOI=3) with RABV or not (Mock). At 16 h p.i., cells were fixed before co-staining with anti-N and anti-PP2A-Aα antibodies and analysis by confocal microscopy. **F)** HEK293T cells were incubated with a medium containing LyoVec alone (Mock) or transfected with 5’ppp-dsRNA using LyoVec (5’ppp-dsRNA). After 6 h, cells were fixed before staining with anti-PP2A-Aα antibody and analyzed by confocal microscopy. **G)** HEK293T cells were co-transfected with plasmids encoding GFP-PP2A-Cα (or EV) and mC-TBK1 or MYC-NAP1 for 24 h. After immunoprecipitation of PP2A-Cα with an anti-GFP antibody, cell lysates (WCL) and immunoprecipitated proteins (IP) were analyzed by WB with anti-mC, anti-GFP, anti-MYC and anti-Tubulin antibodies. **H)** HEK293T cells were co-transfected with plasmids encoding GFP-PP2A-Cα, mC-NAP1 and FLAG-TBK1 (or EV). After 20 h, cells were analyzed by confocal microscopy. On microscopy images, square boxes have been magnified (3X) at the top left. Scale bars: 10µm and 2 µm for insets and FRAP experiments.

**Fig. S7.**
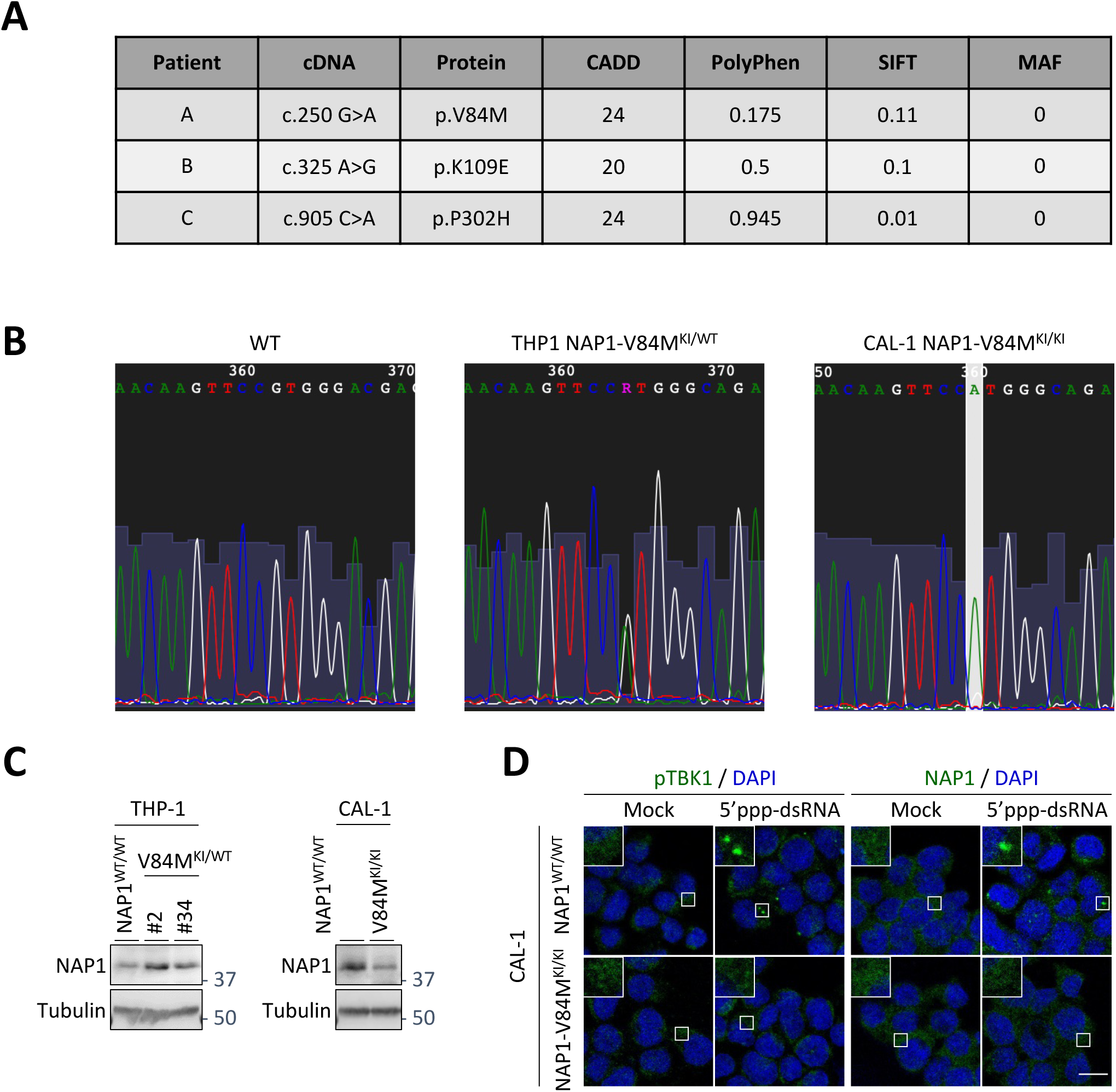
NAP1 V84M variant does not form condensates in CAL-1 cells exposed to a danger signal. **A)** Table showing the characteristics of the different *AZI2*/NAP1 variants. The median allele frequency (MAF) in the general population based on gnomAD database (https://gnomad.broadinstitute.org/), the Combined Annotation Dependent Depletion score (CADD), the Sorting Intolerant from Intolerant score (SIFT), and the PolyPhen score of the different variants are indicated. **B)** Genome sequence electropherograms of THP-1 NAP1^WT/WT^, THP-1 NAP1 V84M^KI/WT^, and CAL-1 NAP1 V84M^KI/KI^ cells showing the heterozygous status of THP-1 NAP1 V84M^KI/WT^ and the homozygous status of CAL-1 NAP1 V84M^KI/KI^ cells. **C)** THP-1 NAP1^WT/WT^, THP-1 NAP1 V84M^KI/WT^, CAL-1 NAP1^WT/WT^, and CAL-1 NAP1 V84M^KI/KI^ cells were lysed. Extracts were analyzed by WB with anti-NAP1 and anti-Tubulin antibodies. **D)** CAL-1 NAP1^WT/WT^ and CAL-1 NAP1 V84M^KI/KI^ cells were incubated with a medium containing LyoVec alone (Mock) or transfected with 5’ppp-dsRNA using LyoVec (5’ppp-dsRNA). After 6 h, cells were fixed before staining with anti-pTBK1 or anti-NAP1 antibody and analyzed by confocal microscopy. On microscopy images, square boxes have been magnified (3X) at the top left. Scale bars: 10µm and 2 µm for insets.

**Fig. S8.**
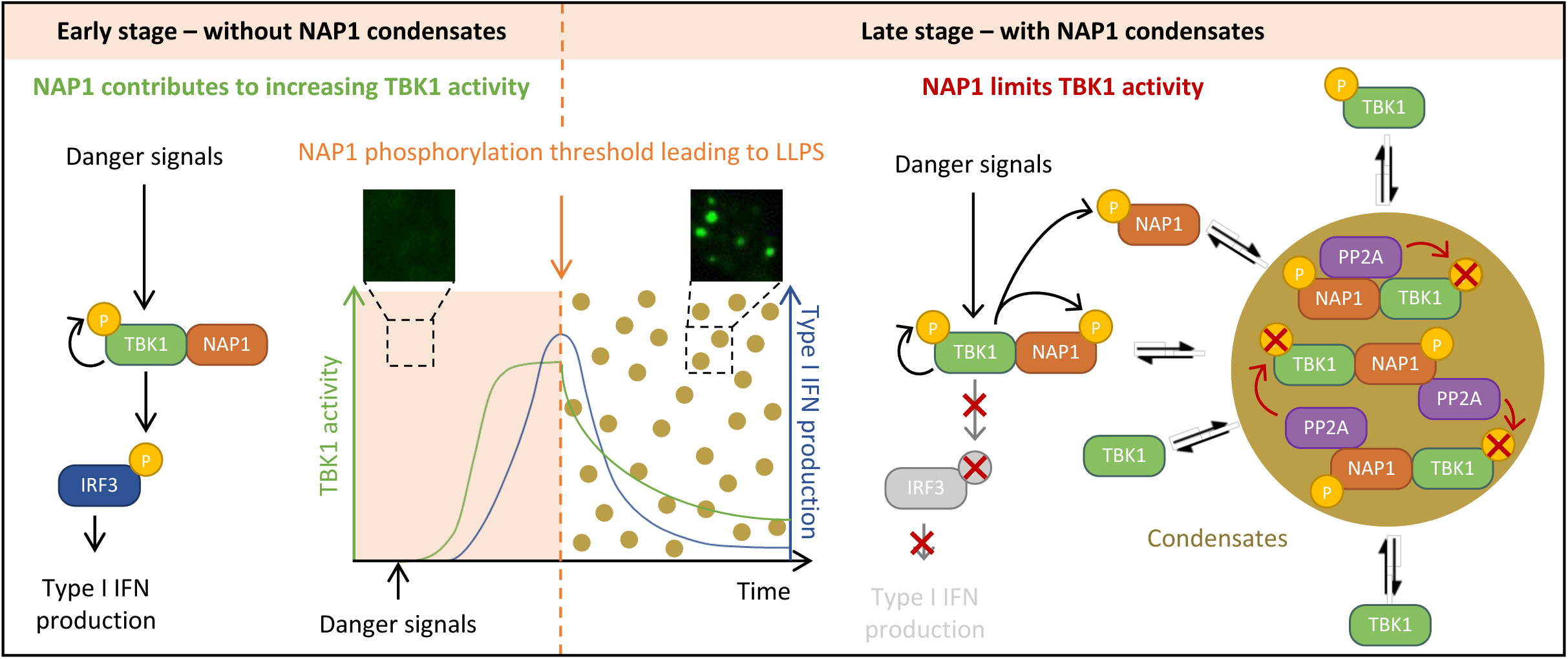
A model for the dual role of NAP1 in the regulation of the IFN-β induction pathway. Upon cellular exposure to danger signals, NAP1 contributes to the activation of TBK1. Being a substrate of the kinase, NAP1 is also phosphorylated. When the amount of phosphorylated NAP1 reaches a critical threshold, it triggers the condensation of the adaptor protein, in which TBK1 and PP2A are concentrated, leading to the dephosphorylation of the kinase and its deactivation.

## Supplementary information

### Case reports

#### Patient A (AZI2: p.V84M)

Female patient followed for an undiagnosed suspected monogenic interferonopathy in France. The clinical presentation is polymorphic and multisystemic, with neurologic, gastrointestinal, pulmonary, hematologic, and immunologic involvement from early childhood.

Neurologically, the patient presents with longstanding spastic diplegia and a pyramidal syndrome, with brain MRI showing diffuse FLAIR and T2 hyperintensities in cortical white matter—predominantly frontal and temporal—as well as calcifications of the basal ganglia. In 2023, she developed a Devic-type neuromyelitis optica (NMO) associated with anti-AQP4 antibodies, presenting with headaches, photophobia. MRI confirmed panmyelitis, and CSF analysis showed hyperproteinorachia and hypoglycorachia without pleiocytosis. After the two first infusions of rituximab, associated with corticosteroid pulses the patient improved. B cells depletion was persistent at 1-year follow-up.

She has a long-standing profile of positive antinuclear antibodies, anti-DNA, and triple antiphospholipid positivity (anti-cardiolipin, anti-b2GP1 and lupus anticoagulant), as well as transient hemolytic anemia episodes, the latest of which occurred in 2022. Lymphocyte phenotyping showed no major quantitative deficits at followed by a persistent B-cell depletion post rituximab as well as a CD4+ and CD8+ T cell lymphopenia. High interferon levels have been consistently recorded since 2018 as well as high interferon signatures.

Pulmonary involvement includes an interstitial lung disease first documented in 2018, with progressive restriction on pulmonary function tests (CPT 69–76%) and a stable decrease in DLCO. Despite this, the patient remains clinically compensated with no oxygen requirement or desaturation on effort. Imaging has shown non-progressive lesions and no active infection, although past pneumococcal pneumonia and arthritis were documented. Current management includes close functional monitoring.

The first symptoms in early life were a transient lacerated gastritis, mild duodenitis and hepatitis. Liver biopsy showed cytolytic hepatitis with lymphocytic infiltration and minimal fibrosis. Hepatic elastography remains normal.

The therapeutic history includes multiple lines: azathioprine and hydroxychloroquine, prolonged corticosteroids, mycophenolate mofetil, aspirin with interruptions due to hematomas, and rituximab. Given the complex autoimmune and inflammatory profile, maintenance treatment with low-dose corticosteroids is currently pursued.

Despite multisystem involvement, the patient maintain stable pulmonary function and neurological status. The overall evolution is consistent with a likely monogenic interferonopathy.

#### Patient B (AZI2 : p.K109E)

This female patient presented with idiopathic thrombocytopenic purpura (ITP) at the age of 3 years, which required corticosteroid therapy due to the ineffectiveness of immunoglobulins. The etiological workup revealed positive antinuclear antibodies (ANA) with a homogeneous and speckled pattern at a titer of 1:2500, while anti-DNA and anti-ENA antibodies were negative.

Six months later, she was tested positive for IgG anticardiolipin antibodies (49 UI/l) and anti-beta2 glycoprotein I antibodies (48 UI/l), without the presence of lupus anticoagulant. Annual follow-ups continued to show positive ANA (titer 1:5000) and elevated levels of IgG anticardiolipin (56 UI/ml) and anti-beta2 glycoprotein I antibodies (61 UI/l), with normal complement levels and no renal involvement.

At the age of 5, following sun exposure, she developed a malar erythematous rash, potentially indicative of discoid lupus, which resolved favorably. She also reported migratory joint pain affecting the cervical spine, upper limbs, knees, and ankles, predominantly in the morning and sometimes causing insomnia, but without any physical limitations or arthritis. Hydroxychloroquine was started together with aspirin.

In the absence of new symptoms or relapse, aspirin and hydroxychloroquine were discontinued at the ages of 12 and 14 years old, respectively.

Autoimmune markers normalized except for persistently positive ANA without specificity and the absence of symptoms suggestive of a connective tissue disease flare.

At the age of 17 years old she is now asymptomatic and off therapy.

#### Patient C (AZI2: p.P302H)

Female patient affected by Down syndrome who developed a complex autoimmune profile combining systemic lupus erythematosus (SLE), type I autoimmune hepatitis (AIH), and intestinal autoimmunity. At age 13, the patient presented with pancytopenia. ANA were strongly positive (>1:1280, speckled pattern), with low-titer anti-dsDNA antibodies. Bone marrow analysis was normal, and macrophage activation syndrome was ruled out. High-dose corticosteroid therapy (1 mg/kg) and hydroxychloroquine were initiated based on the probable diagnosis of SLE, leading to partial hematologic improvement.

Corticosteroid therapy was complicated by induced diabetes managed by insulinotherapy.

At age 14, the patient was diagnosed with type I AIH at the cirrhotic stage. Liver biopsy revealed portal fibrosis, lymphoplasmacytic infiltrates, hepatocellular disorganization, and features consistent with AIH. Autoimmune serology showed positive anti-actin (smooth muscle) antibodies (1:100), ANA (1:800, speckled and homogeneous) and pANCA positivity against MPO. Imaging (MRI, 2016) was consistent with liver fibrosis with early signs of portal hypertension.

Endoscopic evaluations between ages 14 and 17 revealed progressive esophageal varices and portal hypertensive gastropathy, requiring several band ligation procedures. Follow-up imaging at age 16 showed liver dysmorphia with increased splenomegaly and signs of collateral circulation. The last abdominal ultrasound at age 17 confirmed chronic liver changes without suspicious nodules.

Probable autoimmune thyroiditis was suspected during follow-up, though thyroid function tests remained normal, including the last panel at age 17. Two colonoscopies (ages 16 and 17) showed lymphocytic colitis and increased polytypic plasma cells, consistent with autoimmune intestinal involvement.

The therapeutic course included corticosteroids (initiated at age 13 and tapered off by age 17), hydroxychloroquine (stopped at age 14), and azathioprine. This case illustrates a striking constellation of autoimmune disorders—lupus, autoimmune hepatitis and intestinal autoimmunity.

## Notes

### Competing Interest Statement

The authors have declared no competing interest.

